# IQGAP2 regulates blood-brain barrier immune dynamics

**DOI:** 10.1101/2023.02.07.527394

**Authors:** Ketaki A. Katdare, Andrew Kjar, Natasha M. O’Brown, Emma H. Neal, Alexander G. Sorets, Alena Shostak, Wilber Romero-Fernandez, Alexander J. Kwiatkowski, Kate Mlouk, Hyosung Kim, Rebecca P. Cowell, Katrina R. Schwensen, Kensley B. Horner, John T. Wilson, Matthew S. Schrag, Sean G. Megason, Ethan S. Lippmann

**Affiliations:** Vanderbilt Brain Institute, Vanderbilt University, Nashville, TN, USA; Department of Biomedical Engineering, Vanderbilt University, Nashville, TN, USA; Department of Systems Biology, Harvard Medical School, Boston, MA, USA; Department of Chemical and Biomolecular Engineering, Vanderbilt University, Nashville, TN, USA; Department of Neurology, Vanderbilt University Medical Center, Nashville, TN, USA; Vanderbilt Memory and Alzheimer’s Center, Vanderbilt University Medical Center, Nashville, TN, USA

## Abstract

Brain endothelial cells (BECs) play an important role in maintaining central nervous system (CNS) homeostasis through blood-brain barrier (BBB) functions. BECs express low baseline levels of adhesion receptors, which limits entry of leukocytes. However, the molecular mediators governing this phenotype remain mostly unclear. Here, we explored how infiltration of immune cells across the BBB is influenced by the scaffold protein IQ motif containing GTPase activating protein 2 (IQGAP2). In mice and zebrafish, we demonstrate that loss of Iqgap2 increases infiltration of peripheral leukocytes into the CNS under homeostatic and inflammatory conditions. Using single-cell RNA sequencing and immunohistology, we further show that BECs from mice lacking Iqgap2 exhibit a profound inflammatory signature, including extensive upregulation of adhesion receptors and antigen-processing machinery. Human tissue analyses also reveal that Alzheimer’s disease is associated with reduced hippocampal IQGAP2. Overall, our results implicate IQGAP2 as an essential regulator of BBB immune privilege and immune cell entry into the CNS.

## Introduction

Maintenance of central nervous system (CNS) homeostasis is crucial for ensuring normal functions in neurons and glial cells, which are sensitive to exogenous molecules in circulation (*1*). The brain is insulated from these factors with the help of a specialized partition known as the blood-brain barrier (BBB). The BBB is composed of barrier-forming brain endothelial cells (BECs) with unique cellular machinery that regulate the entry of macromolecules and solutes into the brain (*2*), including transporters that control the bidirectional exchange of nutrients and waste and tight junctions that prevent passive leakage of blood components into the brain (*3*).

In addition to modulating molecular transport, BECs also act as a selective interface between the peripheral immune system and the brain (*4*). Until recently, the CNS was considered to be completely isolated from the peripheral immune system and therefore an immune-privileged organ (*5*, *6*). However, recent evidence shows that the CNS is under constant immune surveillance to identify and resolve mediators of injury (*7*, *8*). For example, microglia are tissue resident innate immune cells that continually inspect the CNS parenchyma (*9*, *10*). Further, interstitial fluid and cerebrospinal fluid provide drainage pathways for prospective antigens to reach the periphery and stimulate immune cells (*11–14*). As such, initiation of an inflammatory response in the CNS can lead to recruitment of leukocytes across the BBB. BECs facilitate this process through the expression of various receptors, such as leukocyte adhesion molecules (LAMs), that allow interactions with and extravasation of leukocytes into tissue beds (*15*, *16*). In addition, BEC chemokine signaling and antigen presentation (*17–19*) also play a key role in orchestrating leukocyte extravasation. While the underlying molecular mechanisms of leukocyte extravasation are similar across all organs, BECs have been shown to express very low levels of LAMs under homeostatic conditions, making them refractory to mild inflammatory cues (*1*, *20*). However, despite several exquisite single-cell RNA sequencing (scRNA-seq) studies identifying differentially expressed genes between BECs and peripheral endothelial cells (*21–26*), as well as scRNA-seq profiles of different cell types in the neurovascular unit (*24*, *25*), the mechanisms underlying suppression of LAM expression in BECs and general limitation of immune cell extravasation across the BBB remain poorly understood.

IQ motif containing GTPase activating protein 1 (IQGAP1), a ubiquitously expressed scaffolding protein, has been recently implicated in facilitating leukocyte trafficking across peripheral endothelium (*27*, *28*). Historically, IQGAP1 was known as a regulator of cellular signaling due to its role as a scaffolding protein (*29*). It also acts as an oncogene driving hepatocellular carcinogenesis (*30*, *31*). Interestingly, IQ motif containing GTPase activating protein 2 ( IQGAP2), a related member of the same scaffolding protein family, acts as a tumor suppressor to counteract the oncogenic effects of IQGAP1 (*32*, *33*). Since IQGAP2 is believed to suppress IQGAP1 function, and IQGAP2 expression is predicted in multiple neurovascular support cells such as astrocytes and microglia (*21*, *34–36*), we hypothesized that IQGAP2 may influence inflammatory responses and leukocyte extravasation at the BBB. Herein, using multiple *in vivo* models, we show that loss of Iqgap2 increases leukocyte infiltration into the CNS under various conditions. scRNA-seq of BECs from wildtype and Iqgap2^-/-^ mice further reveals an upregulation of multiple inflammatory genes and signaling pathways involved in BBB-immune cell interactions. Further, using postmortem human brain tissue, we determined that IQGAP2 was reduced in the hippocampus of patients with AD. Overall, our results benchmark IQGAP2 as a key molecular player involved in BBB-immune crosstalk and leukocyte entry into the CNS.

## Results

### Global loss of Iqgap2 increases peripheral immune cell infiltration into the brain in a mouse model of acute neuroinflammation

To initially assess whether murine Iqgap2 influences leukocyte infiltration into the brain, we delivered interleukin 1-beta (IL1β) into the lateral ventricles of wildtype and Iqgap2^-/-^ mice (129S background) to induce acute neuroinflammation (*37*, *38*). After 24 hours, infiltration of leukocytes was assessed by immunohistochemical labeling of CD45+ cells in cortical brain sections. We measured a significant increase in the number of CD45+ cells in the cortex of Iqgap2^-/-^ mice as compared to their wildtype littermates (Figure 1). We also confirmed that delivery of saline to the lateral ventricles in a similar fashion did not stimulate CD45+ cell infiltration into the brains of both wildtype and Iqgap2^-/-^ mice (Figure 1B). These data suggest that Iqgap2 constrains leukocyte infiltration into the mouse CNS following a central inflammatory challenge.

**Figure 1:**
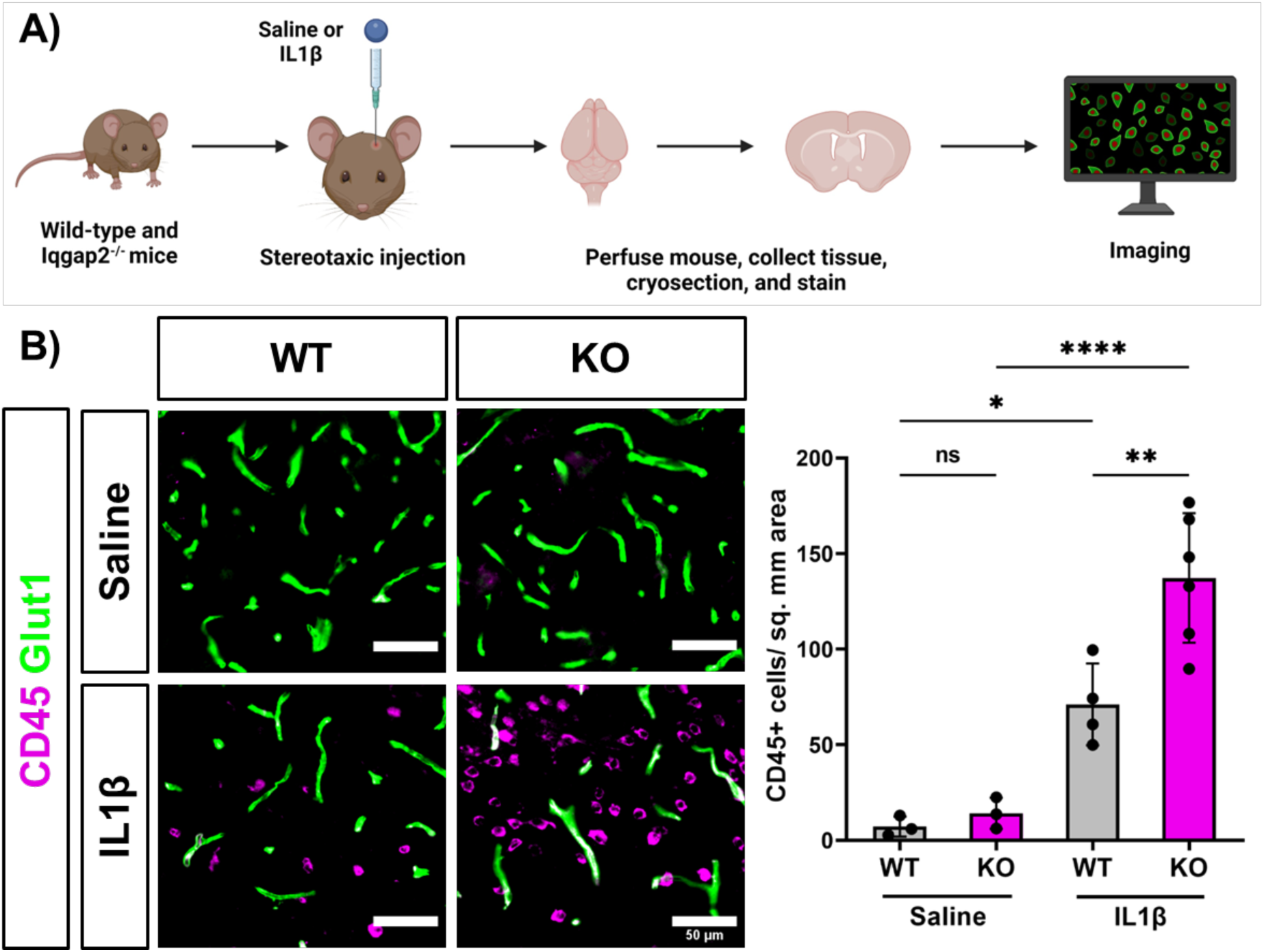
Global loss of Iqgap2 increases infiltration of peripheral leukocytes in a mouse model of acute neuroinflammation. A) Schematic representation of experimental design for assessing leukocyte numbers in mouse brains 24 hours after intracerebroventricular saline or IL1β delivery. B) Representative images and quantification of CD45+ immune cells (magenta) and vasculature (green) in wildtype (WT) and Iqgap2^-/-^ (KO) mouse brain cortex following treatment with saline or IL1β. Data are represented as mean ± SD, and each data point represents an individual mouse, where at least 5 images were quantified per mouse. For saline treatment, N=3 WT mice and N=3 KO mice, and for IL1β treatment, N=4 WT mice and N=6 KO mice. Statistical significance was calculated using one-way ANOVA with Tukey’s multiple comparison’s test (*, p<0.05, **, p<0.01, ****p<0.0001).

### Global loss of Iqgap2 increases immune cell infiltration in experimental autoimmune encephalomyelitis

As dysregulated immune cell infiltration is a hallmark of several neurodegenerative conditions (*39–45*), we sought to next evaluate whether loss of Iqgap2 affects immune cell access to the CNS in the presence of an inflammatory neurodegenerative condition. As such, to monitor the effects of Iqgap2 loss under a chronic inflammatory insult (*46*), we induced experimental autoimmune encephalomyelitis (EAE) in wildtype and Iqgap2^-/-^ 129S mice (Figure 2A). To validate our experimental strategy, we concurrently induced EAE in C57BL/6 mice and observed robust development of disease (Supplementary Figure 1). We measured a significant increase in CD45+ cells in the Iqgap2^-/-^ lumbar spinal cord compared to wildtype 129S mice 30 days after EAE induction (Figure 2B). Interestingly, this increase in infiltrating leukocytes did not produce significant differences in disease severity, probability of survival, or demyelination (Figure 2C-E), suggesting the infiltration events may be decoupled from pathology. Overall, these data further indicate that loss of Iqgap2 contributes to increased leukocyte extravasation into the CNS under extended neuroinflammation.

**Figure 2:**
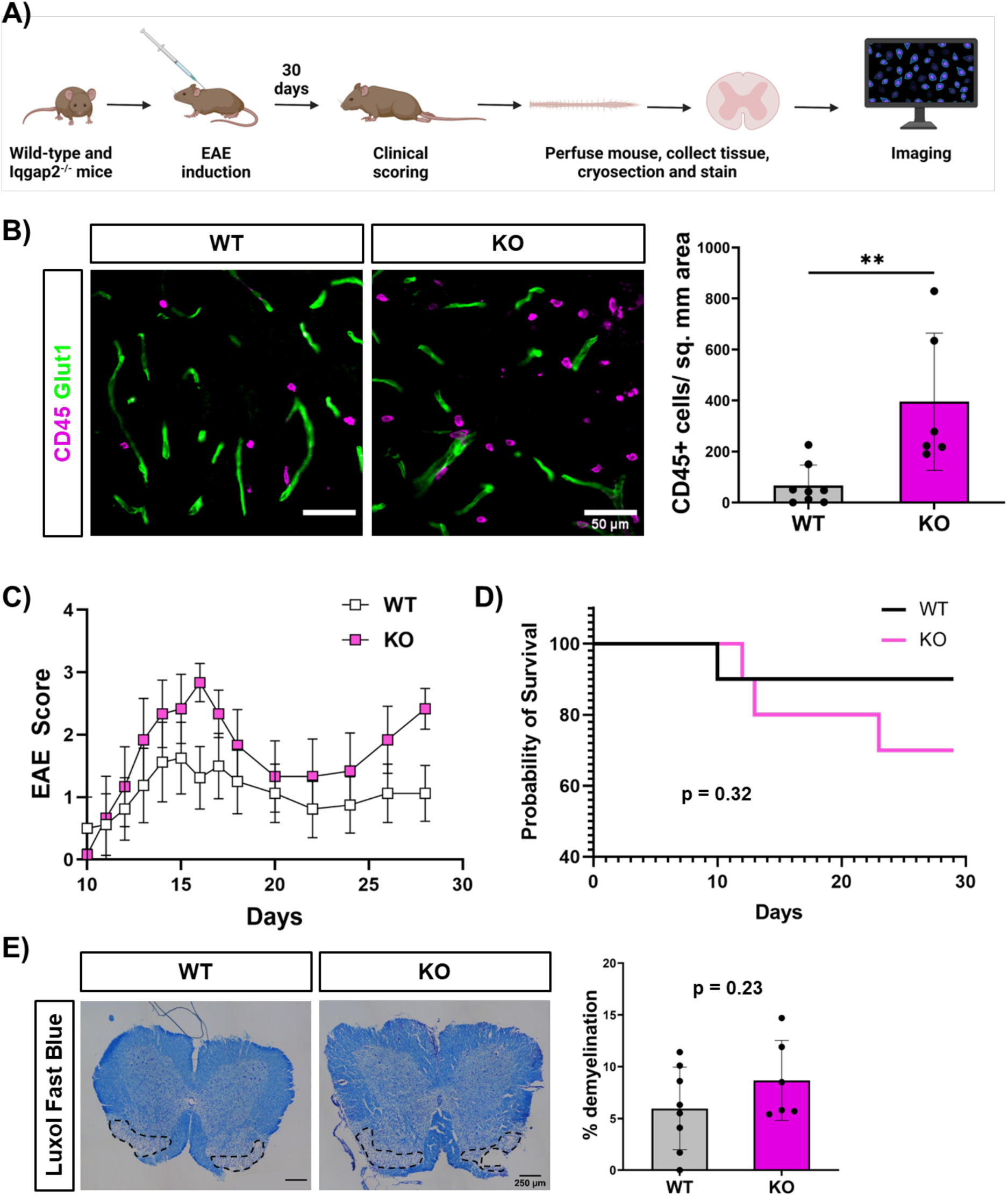
Global loss of Iqgap2 increases infiltration of peripheral leukocytes in EAE. A) Schematic representation of experimental design for assessing response to EAE 30 days following induction. B) Representative images and quantification of CD45+ immune cells (magenta) and vasculature (green) in wildtype (WT) and Iqgap2^-/-^ (KO) lumbar spinal cord at 30 days following EAE induction. Data are represented as mean ± SD, and each data point represents an individual mouse. At least 8 images were quantified per mouse. N=8 WT mice, N=6 KO mice. Statistical significance was calculated using the unpaired student’s t-test (**, p<0.01). C) EAE score curve for wildtype (WT) and Iqgap2^-/-^ (KO) mice following EAE induction. Data are presented as mean ± SEM. N=8 WT mice, N=6 KO mice. Statistical significance was calculated using the unpaired student’s t-test on area under the curve. D) Probability of survival in WT versus KO mice following EAE induction. N=10 WT and KO mice. Statistical significance was calculated using Log-rank (Mantel-Cox) test. E) Representative images and quantification of demyelinating lesions in WT and KO lumbar spinal cord section stained with Luxol Fast Blue at 30 days following EAE induction. Data are represented as mean ± SD, and each data point represents an individual mouse, where 1 image was quantified per mouse. N=8 WT mice, N=6 KO mice. Statistical significance was calculated using the unpaired student’s t-test.

### Loss of Iqgap2 increases infiltration of peripheral immune cells into the brain in zebrafish in the absence of inflammation

Our data in multiple inflammatory mouse models suggest that Iqgap2 plays an important role in BBB immune privilege. To corroborate these findings in an additional species, we generated zebrafish crispants by direct injection of Cas9 protein with multiple sgRNAs to target genes of interest in single-cell embryos with both endothelial cells (kdrl:mCherry) and immune cells (mpeg1:EGFP) transgenically labeled. We specifically assessed the presence of mpeg+ macrophage lineage cells in the brains of 5 days post fertilization (dpf) zebrafish, which have a functional BBB (*47*), that were either uninjected or targeting *tyr* or *iqgap2.* While uninjected fish are expected to retain normal function of all genes, *tyr* crispants should have mosaic knockout of tyrosinase, a protein involved in pigment production that is not expected to affect leukocyte infiltration; this serves as an additional CRISPR injection control. Both controls displayed similarly low numbers of mpeg+ cells in the brain, while *iqgap2* crispants displayed a significant increase in the number of mpeg+ cells (Figure 3B). Since mpeg also labels brain resident microglia, we used an established method to label microglia with Neutral Red dye (*48*) and distinguish these cells from infiltrating leukocytes. Uninjected controls and *iqgap2* crispants were therefore treated with Neutral Red and all mpeg+/Neutral Red+ double-positive microglia and mpeg+/Neutral Red-infiltrating leukocytes were quantified throughout the entire zebrafish brain. We measured a significant increase in the total number of mpeg+ cells, similar to previous experiments, but not double-positive microglia in the *iqgap2* crispants, and the number of mpeg+/Neutral Red-infiltrating leukocytes was significantly increased (Figure 3C). These data suggest that *iqgap2* is essential for restricting the infiltration of leukocytes into the CNS under homeostatic conditions in zebrafish, as its mosaic depletion enhanced the entry of peripheral leukocytes into the brain.

**Figure 3:**
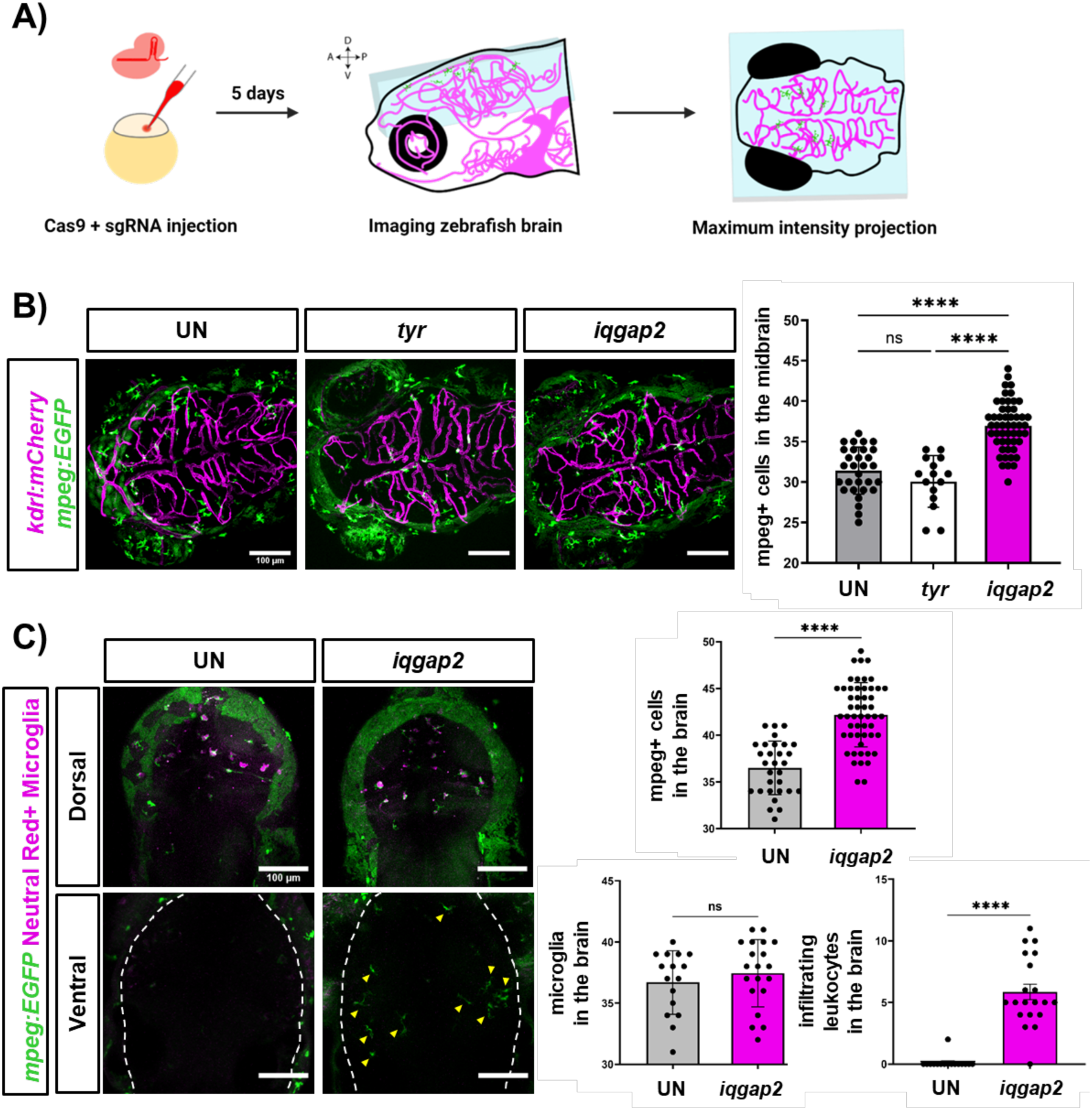
Mosaic loss of *iqgap2* expression increases infiltration of peripheral immune cells into the zebrafish brain. A) Schematic representation of experimental design for assessing leukocyte numbers in the larval zebrafish brain. Double transgenic (kdrl:mCherry; mpeg:EGFP) single-cell embryos were injected with Cas9 protein and sgRNAs to target genes of interest. These mosaic crispants were then allowed to develop normally, and mpeg+ leukocytes were quantified in the brain at 5 dpf. B) Representative 100 µm thick maximum intensity projection images and quantification of macrophage lineage cells (mpeg:EGFP) in the brains of uninjected (UN) controls, *tyr* crispant controls, and *iqgap2* crispants. Vasculature is marked with the kdrl:mCherry transgene (magenta). N=30 fish (UN), 15 fish (*tyr*), and 52 fish (*iqgap2*). Data are represented as mean ± SD, and each data point represents an individual fish. Statistical significance was calculated using a one-way ANOVA with Tukey’s multiple comparison’s test (****, p<0.0001). C) Representative images and quantification of infiltrating leukocytes versus tissue-resident microglia in the brains of *iqgap2* crispants versus uninjected (UN) controls. Representative 30 µm thick maximum intensity projection images of the dorsal (top) or ventral (bottom) brain regions and quantification of mpeg+ (green) macrophage lineage cells and mpeg+/Neutral Red+ (magenta) microglia. Yellow arrowheads indicate individual mpeg+/Neutral Red-infiltrating leukocytes in the ventral brain of *iqgap2* crispants. N=16 fish (UN) and 20 fish (*iqgap2*). Data represented as mean ± SD, and each data point represents an individual fish. Statistical significance was calculated using an unpaired Student’s t-test (****, p<0.0001).

### Global loss of Iqgap2 yields a profound inflammatory transcriptomic profile in mouse BECs

Our data indicate that Iqgap2 plays an important role in restricting peripheral immune access to the CNS and in modulating responses to inflammatory insults. Iqgap2 is a large scaffolding protein known to orchestrate many different cellular functions such as regulating cytoskeletal organization, cytokinesis, and carcinogenesis (*29*), suggesting it could govern many different signaling axes that would influence BBB function and cellular crosstalk within the neurovascular unit. After confirming general expression of Iqgap2 in mouse brain and enriched vessel fractions (Supplementary Figure 2), we performed scRNA-seq on BECs isolated from wildtype and Iqgap2^-/-^ mice to better understand how loss of Iqgap2 affects BBB function. Here, to generate an endothelial cell-enriched population for sequencing, we isolated antibody-labeled CD31+ cells from dissociated mouse brains using fluorescence-activated cell sorting (Figure 4A). After implementing quality control metrics, cells were first analyzed using dimension-reduction by uniform manifold approximation and projection (UMAP) and unsupervised clustering to obtain 12 unique identities: endothelial cells (EC), PLVAP-expressing endothelial cells (EC_plvap), hemoglobin-expressing endothelial cells (EC_hb), endothelial/stromal cells or pericyte-like cells (EC/PC), endothelial/stromal cells or astrocyte-like cells (EC/AC), astrocytes (AC), B cells, T cells, monocytes (MNC), microglia (MG), oligodendrocytes (OLG), and fibroblasts (Supplementary Figure 3). Endothelial cells were the largest represented cell type, followed by immune cells such as monocytes and T cells that are also predicted to express CD31 (Figure 4B and Supplementary Figure 3); other cell types likely represent some small contamination in the sorting process. Each cell type was annotated using previously established marker genes (*49*) and all non-endothelial clusters were filtered out of the dataset for these initial analyses. Wildtype (WT) and Iqgap2^-/-^ (KO) genotypes were equally represented in the EC cluster (Figure 4C), and bulk gene expression comparisons between WT and KO BECs show 928 differentially regulated genes (DEGs) (Figure 4D). Since leukocyte extravasation occurs predominantly at post-capillary venules (*50*–*52*), we examined these DEGs along the neurovascular tree. We used unsupervised clustering to subcluster BECs into 6 sub-populations, and based on marker gene expression, these subclusters were further classified into arterial (A; genes *Hey1, Bmx, and Sema3g*), capillary (C; genes *Slc16a2, Car4, and Mfsd2a*), and venous (V; genes *Icam1, Slc38a5, and Vwf*) zonal identities (Figure 4E and Supplementary Figure 3) (*21*). Further analysis suggests that the strongest DEGs are shared among zonal identities and include genes involved in antigen presentation, interleukin receptor subunits, and adhesion molecules. Unique DEGs identified in the A and V zonal identities did not show any significant functional enrichment whereas a similar analysis of unique DEGs identified in the C zonal identity suggest subtle changes in BEC function (Figure 4F and Supplementary Table 1).

**Figure 4:**
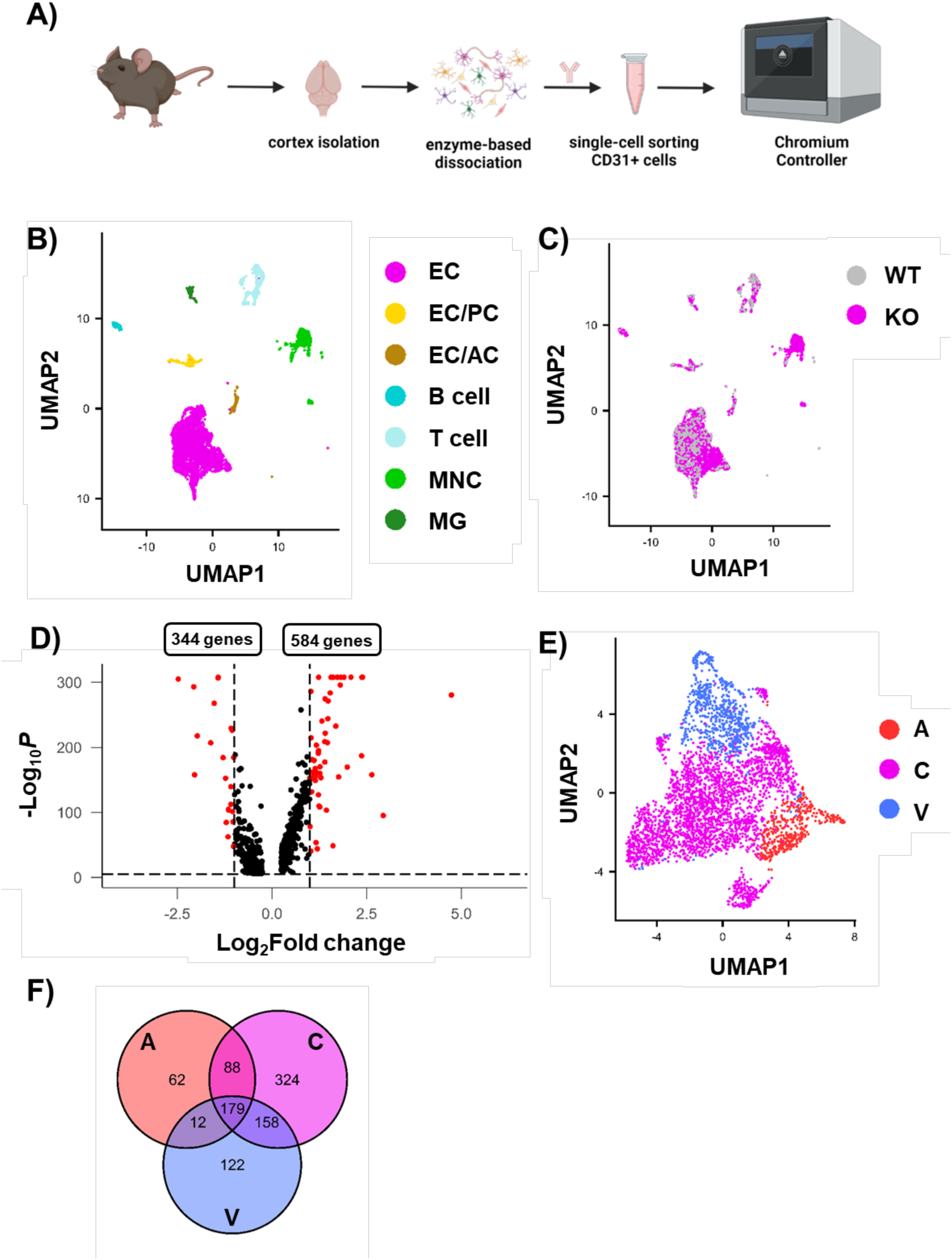
Global loss of Iqgap2 yields extensive transcriptional changes in BECs. A) Schematic representation of experimental design. Whole brain cortices were isolated from wildtype (WT) and Iqgap2^-/-^ (KO) mice, dissociated to a single cell suspension using enzyme-based dissociation techniques, and sorted to enrich for CD31+ cells before sequencing. B) UMAP cell annotations based on unsupervised clustering. EC = endothelial cells, EC/PC = EC/stromal cells (pericytes), EC/AC = EC/stromal cells (astrocytes), MNC = monocytes, MG = microglia. C) UMAP comparison between WT (grey) and KO (pink) cells. D) Volcano plot highlighting differentially expressed genes in the EC cluster. 928 genes were significantly altered, with 584 upregulated and 344 downregulated by loss of Iqgap2. Red dots indicate genes with p<0.05 and >2-fold change in expression. E) UMAP of endothelial zonal identity based on unsupervised clustering. A = arterial, C = capillary, V = venous. F) Venn diagram showing number of DEGs shared between zonal identities.

Across these zones, we were able to confirm that loss of Iqgap2 does not significantly affect expression of most canonical BBB genes, including junction proteins and nutrient transporters such as *Cdh5, Cldn5, Ocln, Tjp,* and *Slc2a1* (Supplementary Figure 4A). At the protein level, total vessel density was unchanged, and we found no obvious deficits in expression of claudin-5, occludin, ZO-1, and Glut1 in the Iqgap2^-/-^ mice (Supplementary Figure 5). We did observe significant differences in gene expression for certain transporters that facilitate exchange of amino acids and metabolites across the BBB, such as *Abcb1a*, *Slc7a1*, *Slc7a5*, and *Slc16a1* (Supplementary Figure 4B). In addition, major regulator genes involved in BBB functional development and maintenance like *Mfsd2a* (*53*) and *Ctnnb1* (*54–56*) were significantly downregulated in the Iqgap2^-/-^ BECs, suggesting possible connections to BBB dysfunction (Supplementary Figure 4C). Interestingly, we also saw significant upregulation of several LAMs and chemokine receptors. Although interaction with leukocytes and response to inflammation is primarily facilitated by venous ECs, we found that loss of Iqgap2 significantly upregulates expression of leukocyte receptors and signaling molecules like *Vcam1, Icam1,* and *Ackr1* across multiple vascular zones (Figure 5A). Using immunohistochemistry, we confirmed upregulation of Vcam1 in cortical vasculature of Iqgap2^-/-^ mice (Figure 5B). The widespread expression of vascular Vcam1 in Iqgap2^-/-^ mice was particularly striking, given that Vcam1 expression on BECs has been implicated in brain inflammation and cognitive decline in mice (*57*). We have also recently shown that vascular VCAM-1 expression is significantly increased in Alzheimer’s disease cortex relative to asymptomatic age-matched controls, further highlighting its links to human neurodegeneration (*58*).

**Figure 5:**
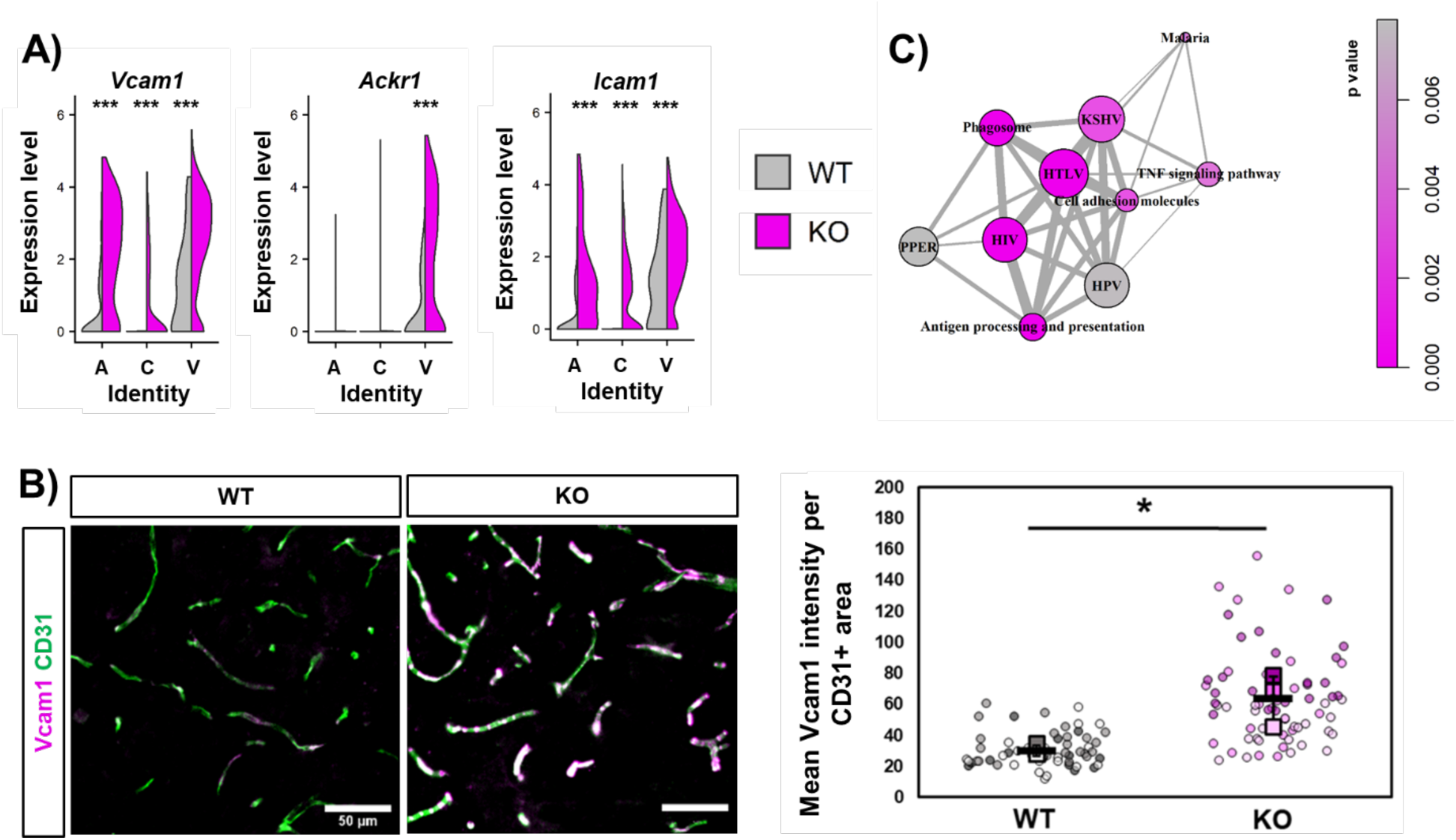
Global loss of Iqgap2 produces a widespread inflammatory phenotype in BECs. A) Split violin plots indicating differential gene expression of select inflammatory markers across vascular zones between wildtype (WT) and Iqgap2^-/-^ (KO) BECs. Statistical significance was calculated using Wilcox rank-order tests with Bonferroni correction (***, p<0.001). B) Representative images and quantification of vascular Vcam1 expression in WT versus KO mouse cortex. Quantification was performed across N=3 biological replicates. Data are represented as mean ± SD (black bars). Biological replicates are represented as squares and measurements from individual images are represented as circles color coded to each replicate. Statistical significance was calculated using the student’s unpaired t-test (*, p<0.05). C) GSEA analysis for signaling pathways upregulated in KO versus WT BECs. Each node represents an enriched gene set belonging to the labeled canonical pathway. Nodes are colored based on p-value, and thickness of the connecting lines indicates similarity of overlapping genes represented in connected gene sets. KSHV = Kaposi sarcoma-associated herpes virus infection, HTLV = Human T-cell leukemia virus 1 infection, HIV = Human immunodeficiency virus 1 infection, PPER = Protein processing in endoplasmic reticulum, HPV = Human papillomavirus infection.

To extend our understanding of potential pathways in BECs affected by global loss of Iqgap2, we performed gene set enrichment analysis (GSEA) for KEGG signaling pathways. GSEA indicated that pathways involved in response to infections like Kaposi sarcoma-associated herpes virus infection (KSHV), human T-cell leukemia virus 1 infection (HTLV), human immunodeficiency virus 1 infection (HIV), and human papillomavirus infection (HPV) were upregulated. In addition, other pathways facilitating immune interactions like cell adhesion (*Vcam1, Icam1*), antigen processing and presentation (*Psme2, Hspa5, Canx, Calr*), and TNF signaling (*Cxcl1, Csf1, Ptgs2*) were also significantly upregulated in the Iqgap2^-/-^ BECs (Figure 5C and Supplementary Figure 6). These results indicate that *Iqgap2* loss shifts both the transcriptional profile and protein expression of BECs towards an activated, inflammatory state.

Due to the significant upregulation of LAMs in the Iqgap2^-/-^ BECs, we further analyzed cell-cell interactions between BECs and other cell types identified in the scRNA-seq dataset using CellChat (*59*). We found that BEC-immune cell interactions were overrepresented in Iqgap2^-/-^ mice. Quantification of these results suggested an increase in cell-cell interactions between BECs and microglia as well as BECs and peripheral immune cells like monocytes, T cells, and B cells. We then assessed the predicted directionality of these interactions by analyzing known receptor-ligand pairs. BECs were predicted to be the “senders” whereas immune populations, especially monocytes, were the primary “receivers” (Supplementary Figure 7). To understand whether these changes were due to Iqgap2 loss in a specific cell type, we performed DEG analyses in all clusters annotated as immune populations. Monocytes had the highest number of significant DEGs, followed by microglia and T cells (Supplementary Figure 8A). KEGG pathway analysis of DEGs in the monocytes indicated upregulation of pathways such as leukocyte transendothelial migration and regulation of actin cytoskeleton (Supplementary Figure 8B). These data indicate that Iqgap2 may also play an important role in modulating BEC-monocyte communication, which could additionally contribute to the overall inflammatory phenotype observed in the BECs in Iqgap2^-/-^ mice.

### IQGAP2 in postmortem human brain tissue

To putatively assess connections between IQGAP2 and human disease states, we evaluated IQGAP2 protein distribution patterns in postmortem human hippocampal tissue from patients with Alzheimer’s disease (AD) and in cases without AD. Using a custom polyclonal antibody raised against a peptide with selective homology to human IQGAP2, we immunostained and quantified vascular-associated and parenchymal IQGAP2 signal in human hippocampal sections. IQGAP2 staining was strongly detected along blood vessels (identified by collagen expression), with more diffuse and punctate signal observed in the parenchyma (Figure 6). A significant decrease in IQGAP2 levels was found in parenchymal tissue (non-vascular regions) in AD patients (Figure 6). Connections between immune cell entry into the brain and neurodegeneration are becoming increasingly scrutinized (*60*, *61*), highlighting the potential importance of this finding.

**Figure 6:**
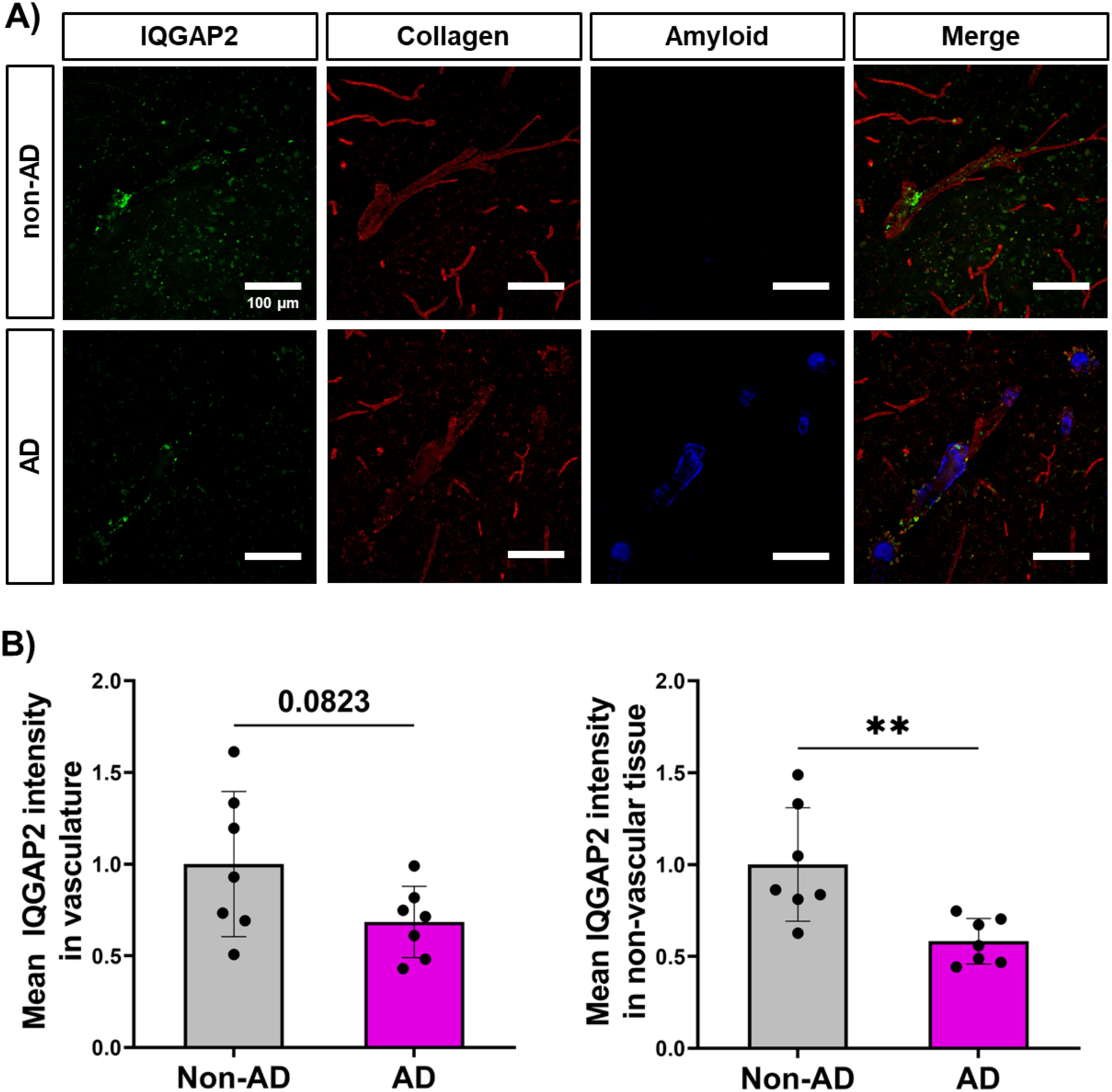
IQGAP2 distribution in human hippocampus. A) Representative confocal microscopy images of IQGAP2 (green) in human hippocampal tissue from AD and non-AD donors. Images were produced from 10 μm z-stack scanning projections with a step interval of 1 μm. Vasculature was stained with Collagen (red) and β-amyloid and neuritic plaques, neurofibrillary tangles and other tau aggregates were stained with Methoxy-X04 (blue). B) Mean IQGAP2 intensity in vascular regions was quantified solely one scanning projection in collagen+ area, while IQGAP2 intensity in parenchyma was quantified by gating samples to exclude collagen+ area. Data are presented as mean ± SD, where each data point represents an individual donor with at least five images quantified per donor. N=7 non-AD donors and N=7 AD donors. Statistical significance was calculated using unpaired student’s t-test (**, p<0.01).

## Discussion

Peripheral endothelial cells display high levels of LAMs and can respond swiftly to local and systemic inflammatory cues (*62*). This is followed by rapid infiltration of leukocytes into surrounding tissue beds. In comparison, BECs are comparably immune quiescent under homeostatic conditions and generally express low levels of LAMs. This allows the BBB to more selectively control the activation of downstream inflammatory pathways and extravasation of leukocytes into the CNS (*63*, *64*). Although peripheral immune responses are essential for the resolution of CNS injury, it is well established that age-related neurological deficits and chronic neurodegeneration may be exacerbated in part by unwarranted entry of leukocytes into the brain (*39*, *41*, *65–67*). Further, BECs upregulate transcriptional signatures of inflammatory responses during aging and neurological disease (*24*, *25*, *68*). As such, identifying mechanisms that regulate BEC inflammatory responses is critical for understanding the pathological progression of these diseases and developing strategies to decrease leukocyte extravasation. Our study provides evidence that IQGAP2 plays an important role in BBB immune dynamics and the propensity of leukocytes to enter the CNS after immune stimulation.

To our knowledge, the role of IQGAP proteins in BBB integrity has not been studied. *IQGAP2* belongs to the *IQGAP* family of scaffolding proteins involved in orchestrating a wide array of intracellular signaling and cytoskeleton dynamics (*29*). The multidomain structure of these proteins acts as a framework for complex formation of signaling proteins, thus influencing many downstream cellular processes. *IQGAPs* were historically considered to be modulators of cytoskeletal architecture. However, it has become apparent that their role extends into other physiological processes like glomerular filtration in the kidney, cardiomyocyte function in the heart, smooth muscle cell contraction in lung airways, and metabolism in the liver (*29*, *69*, *70*). *IQGAP2* has also been studied in the context of its tumor-suppressive characteristics (*71*). Previous studies identified *IQGAP2* as a novel tumor suppressor gene specifically linked to the development of hepatocellular carcinoma. More recently, *IQGAP2* inactivation has been linked to other malignancies like gastric cancer (*72*), prostate cancer (*73*), and bladder cancer (*74*). Moreover, reduced expression of *IQGAP2* is associated with worsened cancer pathology and poor clinical outcomes (*73*, *75*, *76*). Our study provides new context for the role of IQGAP2 in physiological processes related to immune cell trafficking.

One limitation of our study is that we cannot exclusively ascribe the observed CNS leukocyte infiltration to loss of Iqgap2 expression in a particular cell type*. IQGAP2* has generally high expression in many immune cell subtypes (*77*), but this has not previously been associated with specific phenotypes like tissue extravasation and responsiveness to inflammatory cytokines. Our current data may suggest that cell-to-cell communication between BECs and immune cells plays a role in BBB inflammation when Iqgap2 is lost. However, more extensive scRNA-seq profiling of the neurovascular unit in Iqgap2^-/-^ mice, as well as lineage-specific Iqgap2 knockout models, will be necessary in future studies to fully clarify cell-intrinsic and cell-extrinsic effects of IQGAP2 on BBB immune privilege.

An additional unanswered question is whether changes in IQGAP2 expression contribute to human neurodegenerative disease progression through modulation of immune cell recruitment to the CNS. In the acute and chronic inflammatory animal models used in this study, we did not observe significant differences in pathology. In the EAE model, it is possible that the 30-day timepoint does not reflect increased damage caused by infiltrating leukocytes due to other compensatory mechanisms (e.g. significant damage could occur at an earlier time point followed by regeneration), especially when considering the biphasic disease severity in the Iqgap2^-/-^ mice and that deaths generally occurred during periods where disease severity worsened. In human hippocampal tissue, we observed a decrease in IQGAP2 in patients with AD. Single-cell datasets indicate parenchymal IQGAP2 expression is restricted to microglia, while vascular IQGAP2 expression could potentially be localized to perivascular macrophages or fibroblasts (*21*, *24*, *36*). Interestingly, microglia have been recently connected to T cell infiltration in mouse models of tauopathy (*78*) and general aging (*79*), and T cell infiltration and resultant activation of microglia has been shown to exacerbate neurodegeneration in engineered human cell-based models of AD (*80*). In the context of our study, these findings motivate future exploration into whether loss of Iqgap2 influences pathology and immune cell infiltration in mouse and human models of AD.

Overall, our work reveals a novel role for IQGAP2 in regulating BBB immune dynamics. While the cell-specific effects of IQGAP2 are currently unclear, our collective data suggest that this protein plays an important conserved and previously unrecognized role in suppressing BEC inflammatory responses and regulating immune cell trafficking to the CNS through non-cell autonomous mechanisms. Future work will determine these mechanisms of action and their relevance to brain disorders.

## Methods

### Mouse maintenance and procedures

#### Colony maintenance

All mouse protocols were approved by the Institutional Animal Care and Use Committee at Vanderbilt University. Iqgap2^-/-^ mice (129S background) were obtained from Jackson Laboratory (strain 025452). Male and female Iqgap2^-/-^ mice and wildtype littermate controls were used for all experiments. Mice were at least 8 weeks of age at the time of use and were housed continuously in an environmentally controlled facility in a 12-hour light/dark cycle with *ad libitum* access to food and water.

#### Genotyping

Mice were ear-tagged and tail snips were collected at approximately 2 weeks of age. Genomic DNA was extracted using an Extracta DNA Prep kit for tissue (Quantabio) per manufacturer instructions. DNA was extracted in the Extraction buffer at 95°C for 30 minutes, cooled to room temperature and mixed with Stabilization buffer before being stored at -20°C. The reactions were performed using the Apex Hot Start Taq BLUE Master Mix (Apex Bioresearch) on a ProFlex PCR system (Applied Biosystems). A touchdown cycling protocol was used with an initial annealing temperature of 65°C gradually lowered to 60°C over the course of 10 cycles. Genotyping primers used were as follows: Mutant Reverse– ATTTGTCACGTCCTGCACGACG, Wildtype Reverse–TGGCCTTCCTCCCTTAAAGT, and Common Forward–TGACTCAGAGGGCACATGGT. PCR products were run on a 2% agarose-TAE buffered gel supplemented with SYBR Safe DNA Gel Stain (Invitrogen) and imaged using a LI-COR Odyssey Fc gel imager (Supplementary Figure 9).

#### Tissue collection

Mice were deeply anesthetized using high-dose isoflurane and euthanized by transcardial perfusion of 1X DPBS (Gibco), followed by 4% paraformaldehyde (PFA, Thermo Fisher Scientific). Brains and lumbar segment of spinal cords were extracted and postfixed in 4% PFA overnight followed by cryopreservation in sucrose gradient solutions (15% and 30%, respectively). The tissue was then embedded in OCT medium (Tissue-Tek) and 15 µm (brain) and 25 µm (spinal cord) thick sections were cut and stored at -80⁰C.

#### Microvessel isolation

Mice were deeply anesthetized using high-dose isoflurane and euthanized by decapitation. Brains were extracted and collected in ice-cold PBS. Microvessels from the cortex were isolated as previously described (*81*). In brief, cortices were dissected from remaining brain tissue, homogenized in PBS using a tissue homogenizer (Wheaton), and collected by centrifugation. Homogenized tissue was resuspended in a 15% dextran solution (∼70,000 kDa, Sigma) and centrifuged at 10,000xg to separate the vessel fraction from the remaining tissue. The vessel fraction was washed with PBS and filtered using a 40 µm cell strainer (Corning). For protein extraction, the vessel fraction was incubated in RIPA buffer (Sigma) supplemented with 1% v/v protease and phosphatase inhibitor cocktails (Sigma) for 30-60 minutes on ice. Cell debris was separated by centrifugation (12,000xg for 15 minutes at 4⁰C) and the supernatant was stored at -20⁰C. Protein concentration was quantified using a Pierce BCA Protein Assay (Thermo Fisher Scientific) according to the manufacturer instructions. For immunohistochemical analysis, microvessel suspension was placed on glass slides and allowed to dry at room temperature. Dried microvessels were then fixed with 4% parafolmaldehyde solution and labelled with Lectin DyLight 488 for 30 minutes at room temperature before mounting in Prolong Gold Antifade Mountant (Invitrogen).

#### Intracerebroventricular injection of IL1β

Male and female Iqgap2^-/-^ mice and wildtype littermates were used as experimental animals. All animals were at least 12 weeks old at the time of the surgery. Under isoflurane anesthesia, mice were unilaterally injected into the lateral ventricle using a stereotactic apparatus at coordinates of AP = -0.3 mm, ML = -1 mm, and DV = -3 mm. After injections, mice were returned to prewarmed home cages for recovery. Each mouse received 20 ng/µL of IL1β solution (20 ng in 1 µL PBS) or an equivalent volume of sterile PBS. 24 hours after surgery, mice were transcardially perfused with PBS followed by 4% paraformaldehyde and brains were extracted for immunohistological analysis. For quantification, CD45+ cells were manually counted in each section under blinded conditions. Vasculature was labelled using a fluorescence-conjugated GLUT1 antibody to solely quantify CD45+ cells that had extravasated out of the vessels into the brain parenchyma.

#### Experimental autoimmune encephalomyelitis (EAE)

Female Iqgap2^-/-^ mice and wildtype littermates were used as experimental animals. All animals were between 9 and 13 weeks of age at the time of induction. EAE kits (Hooke Laboratories) targeting MOG35-55 antigen were used. 100 µL of MOG35-55/Complete Freund’s Adjuvant emulsion was injected subcutaneously at the scruff of the neck and near the base of the tail resulting in a total injection volume of 200 µL into each mouse. At 2 and 24 hours post injection of emulsion, 100 µL of pertussis toxin (4 µg/mL) was injected intraperitoneally. Clinical scores were evaluated starting 7 days post induction as follows: score 1, flaccid tail; score 2, weak hind limbs; score 3, hind limb paralysis; score 4, quadriplegia. Clinical scores were recorded every day for the first week after development of symptoms followed by every other day thereafter. Premature deaths were recorded. 30 days following induction, mice were transcardially perfused with PBS followed by 4% paraformaldehyde. Brains and spinal cords were extracted for immunohistological analysis. For quantification, CD45+ cells were manually counted in each section under blinded conditions. Vasculature was labelled using a fluorescence-conjugated GLUT1 antibody to ensure that CD45+ cells had extravasated out of the vessels. For quantifying EAE pathology in spinal cord, total area and demyelination area were calculated using the “Measure” tool in ImageJ by manually outlining regions of interest as indicated by Luxol Fast Blue stain.

### Zebrafish maintenance and procedures

Zebrafish were maintained at 28.5°C following standard protocols (*82*). All zebrafish work was approved by the Harvard Medical Area Standing Committee on Animals under protocol number IS00001263-3. Adult fish were maintained on a standard light-dark cycle from 8 am to 10 pm. Adult fish, aged 3 months to 1.5 years, were crossed to produce embryos and larvae. For imaging live larvae, 0.003% phenylthiourea (PTU) was used beginning at 1 dpf to inhibit melanin production. These studies used the AB wildtype strains and the transgenic reporter strains (Tg(kdrl:HRAS-mCherry)^s896^ (*83*), abbreviated as Tg(kdrl:mCherry), and Tg(mpeg1:EGFP)^gl22^ (*84*), abbreviated as Tg(mpeg1:EGFP). Mosaic *iqgap2* crispants were generated by injection of 7 µM Cas9 protein complexed with four sgRNAs (5’-AGTAGCCTCGATTTACAGG-3’, 5’-GCACTTTGTCAGTCACGGAA-3’, 5’-CAGGACAGCGCGAGCACTG-3’, and 5’-AAAGTCCGCGCGCAGTTTA-3’) to target multiple sites in the *iqgap2* transcript, and *tyr* control crispants were similarly targeted with four sgRNAs (5’-GCCGCACACAGAGCCGTCGC-3’, 5’-GGATGCATTATTACGTGTCC -3’, 5’-GACTCTACATCGGCGGATGT-3’, and 5’-GTATCCGTCGTTGTGTCCGA-3’). To distinguish between microglia and macrophages, 4 dpf larvae were exposed to 2.5 μg/ml of Neutral Red (Millipore Sigma: N7005) in embryo water for 3 hours at 28.5°C. Larvae were washed at least three times to remove the residual dye and then microglia were assessed the next day as previously described (*48*). Zebrafish larvae were immobilized by tricaine exposure and live imaged on a Leica SP8 line scanning confocal microscope. Quantification of mpeg+ and Neutral Red+ cells was manually performed on blinded z-stack images that spanned the entire larval head using ImageJ.

### Human brain tissue and preparation

Human brain tissue was obtained at autopsy and prepared as previously described (*85*). De-identified brain tissue was obtained from the Vanderbilt Brain and Biospecimen Bank at Vanderbilt University Medical Center. Written informed consent for brain donation was obtained from patients or their surrogate decision makers. All brain tissue collection was authorized by the Institutional Review Board at Vanderbilt University Medical Center. Demographics and neuropathological information for each donor are listed in Supplementary Table 2.

Human brain tissue was obtained at autopsy and immersion fixed in 10% formalin (Thermo Fisher Scientific) at 4°C for 1-3 days. The fixative solution was then removed and the tissued rinsed with 1x TBS (Corning) three times for 5 minutes each. The tissue was placed in sterile 10% sucrose (Millipore Sigma) /1x TBS/0.02% sodium azide (NaN3, Millipore Sigma) until tissue sank and then 30% sucrose/1x TBS/0.02% NaN3 for overnight at 4°C or until the tissue sank. The tissue block was affixed to the stage of vibratome using cyanoacrylate cement and fully submerged in 1x TBS. Hippocampal sections were prepared at 50 µm thickness. Floating tissue sections were transferred to 15 mL Falcon tubes with antigen retrieval buffer (10 mM citric acid pH 6.0, Millipore Sigma) containing 0.05% Tween-20 (Millipore Sigma) and heated to 95°C for 20 minutes in a block heater. Hippocampal sections were then washed with 100 mM glycine (Millipore Sigma)/1x TBS/0.1% Triton X-100 (Millipore Sigma) buffer for 30 minutes followed by permeabilization with 0.3% Triton X-100/1x TBS buffer for 30 minutes and two washes for 5 minutes each with 1x TBS at room temperature.

### Development of a custom antibody against human IQGAP2 protein

A peptide corresponding to amino acid residues 1460-1474 of human IQGAP2 (RSIKLDGKGEPKGAK) was synthesized with an amino-terminal cysteine and conjugated to maleimide-activated Keyhole Limpet Haemocyanin (KLH), maleimide-activated bovine serum albumin (BSA), and SulfoLink resin using manufacturer protocols (Thermo Fisher). The peptide-KLH conjugate was used to immunize rabbits (Cocalico Biologicals). Rabbit antisera were tested for the presence of antibodies recognizing the IQGAP2 peptide by dot blot analysis using the peptide-BSA conjugate. The rabbit antibodies were affinity-purified from the antisera using the peptide-SulfoLink resin, where 5 mL of rabbit sera was diluted 1:1 with PBS and passed over a 2 mL peptide-SulfoLink column. After extensive washing with PBS, bound antibodies were eluted with 8.5 mL 0.1 M Glycine (pH 2.2) and collected in a tube containing 1.5 mL of 1 M Tris (pH 8). Antibody solution was stored at -80°C.

### Immunofluorescent staining

#### Mouse tissue

Tissue slices were retrieved from the -80⁰C freezer and allowed to thaw at room temperature for 10-15 minutes. Sections were washed with 1X PBS with 0.03% Triton X-100 to remove the OCT medium. Sections were then blocked using a goat serum blocking buffer and incubated in primary antibody solution overnight at 4°C. After incubation, primary antibody solution was thoroughly washed off and sections were incubated in secondary antibody solution for 2 hours at room temperature. All antibodies and corresponding dilutions used for immunohistochemistry are listed in Supplementary Tables 3-4. Following final washes, tissue was mounted in Prolong Gold Antifade Mountant with DAPI (Invitrogen) and slides were allowed to dry overnight before imaging on a Leica DMi8 epifluorescence microscope. All acquired images were processed and quantified using ImageJ software. For quantification of vascular Vcam1 expression, mean Vcam1 intensity was calculated within CD31+ vessels using ImageJ.

#### Human tissue

Immunohistochemical labeling in hippocampal tissue slices was performed as previously described (*85*, *86*) with minor modifications. Tissue slices were incubated with primary antibodies for 48 hours at 4°C followed by secondary antibodies for 2 hours at room temperature. All antibodies and corresponding dilutions used for immunohistochemistry are listed in Supplementary Tables 3-4. Neuritic plaques, neurofibrillary tangles and related AD pathological structures were additionally stained using 1 µM 4,4’-[(2-methoxy-1,4-phenylene) di-(1*E*)-2,1-ethenediyl] bisphenol (MX-04) (Tocris) for 15 minutes at room temperature. Confocal images were acquired using the Zeiss LSM 710 confocal laser-scanning microscope (Carl Zeiss AG) with a 20× air/dry or 63× oil objective and 10 μm z-stack scanning projections with a step interval of 1 μm or one scanning projection, with a minimum resolution of 1500 x 1500 pixels. Vascular IQGAP2 expression was quantified using mean IQGAP2 intensity within Collagen+ area and parenchymal IQGAP2 expression was quantified by gating Collagen-area using ImageJ.

### Luxol Fast Blue staining

Tissue slices were retrieved from the -80⁰C freezer and allowed to thaw at room temperature for 10-15 minutes. Sections were first allowed to dry overnight at room temperature and then immersed in a 70% ethanol solution overnight to facilitate defatting. Luxol Fast Blue stain (Abcam) was applied to the sections and incubated for 5 to 6 hours at 60⁰C. Excess stain was washed off by consecutive dipping in fresh absolute ethanol. Slides were differentiated briefly using lithium carbonate solution and washed with distilled water and alcohol solution. Lastly, sections were counterstained with Cresyl Etch Violet, washed with distilled water, dehydrated with absolute alcohol and mounted in DPX mounting medium (Sigma-Aldrich). Mounting medium was cured overnight before imaging on a Leica DMi8 inverted microscope.

### Western blotting

Protein samples from mouse tissue were prepared by diluting 20-40 μg of protein with 1X Laemmli buffer (Biorad) supplemented with beta-mercaptoethanol (Sigma) and Ultrapure water (Gibco) to a final volume of 20-30 μL. Samples were then boiled at 95⁰C for 5 minutes, cooled on ice, and centrifuged briefly. Samples were then loaded into 4-20% Criterion TGX Midi protein gels (Biorad) along with Precision Plus Dual Color Protein ladder (Biorad) and run at 80-120V. Protein gels were then transferred onto iBlot2 Nitrocellulose membranes (Thermo Fisher Scientific) using an iBlot2 transfer device (Thermo Fisher Scientific). Membranes were cut to size and blocked for at least 30 minutes at room temperature in Intercept TBS Blocking buffer (LI-COR Biosciences) on a shaker. Membranes were submerged in primary antibodies diluted in blocking buffer with 0.05% Tween20 (Sigma) and incubated at 4⁰C overnight. Following primary antibody incubation, membranes were washed in wash buffer (1X tris buffered saline with 0.05% Tween-20). Membranes were incubated in secondary antibodies diluted in the wash buffer at room temperature for 2 hours. All primary and secondary antibody information is listed in Supplementary Tables 3-4. Blots were imaged using a LI-COR Odyssey Clx or Fc Imager and bands were quantified using Image Studio Lite software.

### Single-cell RNA sequencing

Fresh cortical tissue was homogenized and delipidated using an Adult Brain Dissociation Kit (Miltenyi Biotec) according to the manufacturer’s instructions. The resultant single-cell suspension was incubated with TruStain FcX (Biolegend) for 10 minutes at 4°C to prevent non-specific antibody binding of Fc receptors, labeled with a secondary conjugated CD31 antibody (1:2000, eBioscience) for 30 minutes, and counterstained with DAPI. Live CD31+ cells were flow sorted using a 4-laser FACS Aria III sorter (BD Biosciences) at the Vanderbilt Flow Cytometry Shared Resource. Live cells were resuspended in DMEM (Gibco) supplemented with 2% FBS to obtain a concentration of 700-1200 cells/μL. RNA extraction, 10X Genomics Chromium 5’ scRNAseq Library Prep, and sequencing on an Illumina NovaSeq6000 sequencer was performed at the VANTAGE core facility.

### Single-cell RNA sequencing analysis

All gene by counts data were read into Seurat (v.4). The initial data were filtered to retain only cells with RNA counts between 1,200 and 20,000 (with less than 10% being mitochondrial). Within each biological experiment, data were log normalized. The top 2,000 variable features (as identified by variance) were selected. Samples were combined using CCA (Seurat v.4) and then standardized. UMAP based on the first 50 principal components was used to reduce dimensionality for visualization and clustering (*87*). Predicted doublets were filtered out using DoubletFinder (*88*). Unsupervised clustering was achieved using the KNN-graph approach in Seurat (*89*), and clusters were annotated with SingleR (*90*) based on previously published datasets (*49*). Predominantly endothelial clusters (0,2,4,5) were rescaled and clustered based on the first 15 principal components. Clusters were annotated as arterial, capillary, or venous based on the expression of established marker gene sets (*21*). Approximately 2,000 ECs were analyzed per condition. Differentially expressed genes (DEGs) were computed in Seurat using the FindMarkers() function with built-in Bonferroni correction. KEGG or Reactome gene set enrichment analysis (GSEA) was computed on significant DEGs using WebGestalt with a false discovery rate cutoff of 0.05 and weighted set cover redundancy reduction (*91*). Communication between cell types was predicted using the CellChat R packages (*59*). For visualization, *EnhancedVolcano* (*92*), *UpSetR* shiny app (*93*), *iGraph* (*94*), and *ggvenn* were employed.

### Statistical analysis

Statistical analysis for single-cell RNA sequencing data was conducted in R. All other analysis was performed in GraphPad Prism 9.0.0.

## Data availability

Raw sequencing data files have been deposited in the ArrayExpress collection under accession code E-MTAB-12687. All code used for the single-cell RNA sequencing analyses is publicly available at: https://github.com/LippmannLab/IQGAP2_WT_KO_BEC_scRNAseq.

## Author contributions

KAK, ESL, and EHN conceived the study. KAK and ESL designed the majority of experiments with input from the other authors. KAK, NMO, EHN, AGS, AS, AJK, HK, and RPC conducted all experiments. AK performed all bioinformatics analyses. KM assisted with data quantification. WRF performed human tissue histology. KRS and KBH provided support on animal husbandry and takedowns for experiments. MSS provided human tissue samples and contributed to interpretation of histological images. JTW and SGM provided input on experimental planning and data interpretation.

## Acknowledgments

Funding was provided by a Chan Zuckerberg Initiative Ben Barres Early Career Acceleration Award (grant 2019-191850 to ESL) and NIH grant R21 NS106510 (to ESL). AK and EHN were supported by the National Science Foundation Graduate Research Fellowship Program. NMO was supported by a Damon Runyon Postdoctoral Fellowship. Support for RNA sequencing was provided by Vanderbilt Technologies for Advanced Genomics core facility, which is supported by the CTSA Grant (NIH grant 5UL1 RR024975), the Vanderbilt Ingram Cancer Center (NIH grant P30 CA68485), the Vanderbilt Vision Center (NIH grant P30 EY08126), and NIH/NCRR grant G20 RR030956. Support for histology was provided by Vanderbilt Translational Pathology Shared Resource core facility, which is supported by P30 CA068485. Flow cytometry experiments were performed in the Vanderbilt Flow Cytometry Shared Resource, which is supported by P30 CA068485 and the Vanderbilt Digestive Disease Research Center (P30 DK058404). The custom IQGAP2 antibody in this study was produced by the Vanderbilt Antibody and Protein Resource, which was supported by P30 CA068485. Image acquisition was performed in part through the use of the Vanderbilt Cell Imaging Shared Resource, which is supported by NIH grants P30 CA68485, P30 DK20593, P30 EY08126, and S10RR027396. The authors thank Dr. Jose Gomez for early assistance with the Iqgap2^-/-^ mice and Dr. Eric Shusta for sharing microarray data that initially indicated the potential relevance of IQGAP2. Some figures in this manuscript were created in part using BioRender.

