## Supplemental information for "IQGAP2 regulates blood-brain barrier immune dynamics"

Ketaki A. Katdare *et al.*

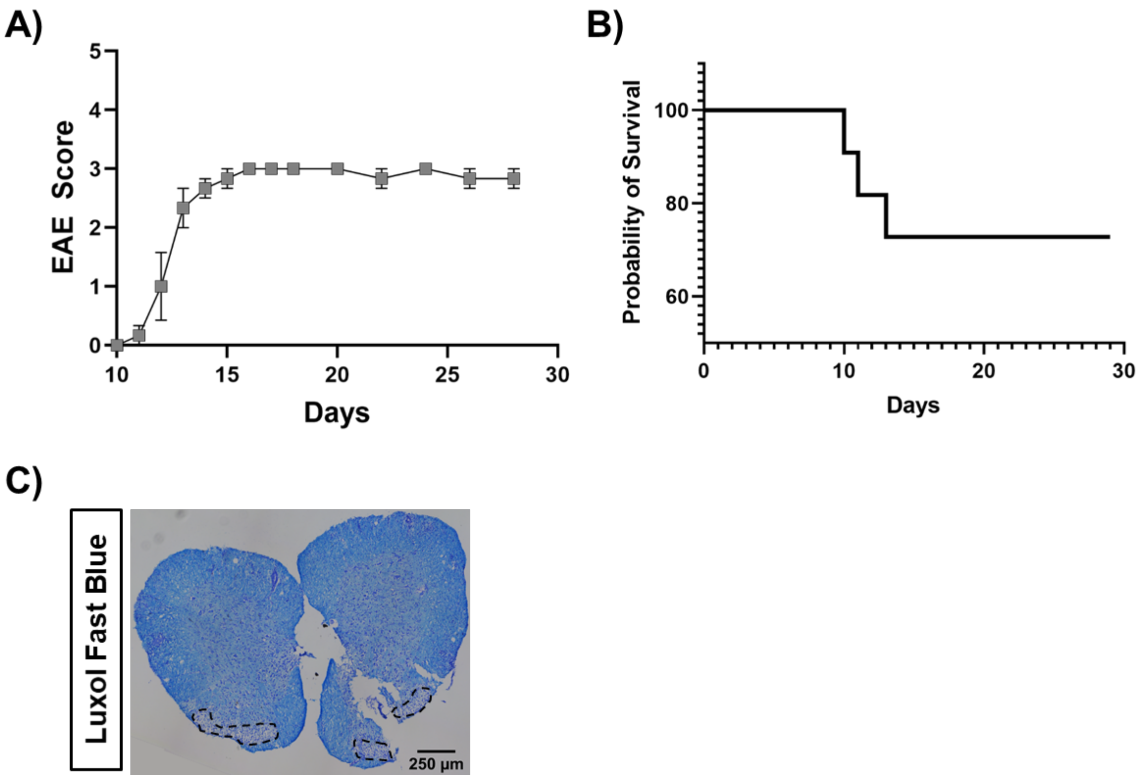

**Supplementary Figure 1: EAE phenotype in C57BL/6 mice.**

A) EAE score curve for C57BL/6 mice following EAE induction. Data are represented as mean ± SEM. N=3 mice.

B) Probability of survival in C57BL/6 mice following EAE induction. N=11 mice.

C) Representative histological images of lumbar spinal cord sections stained with Luxol Fast Blue showing demyelinating lesions. Demyelinated areas are outlined with dotted boundary.

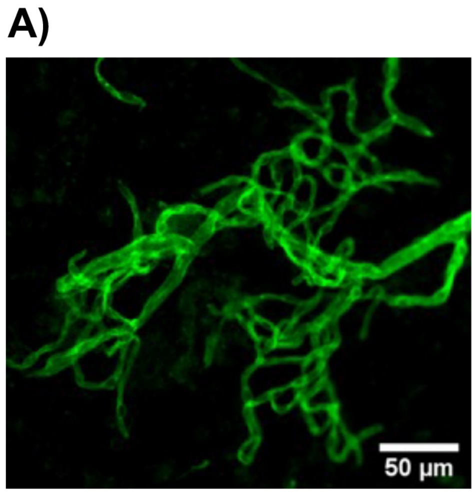

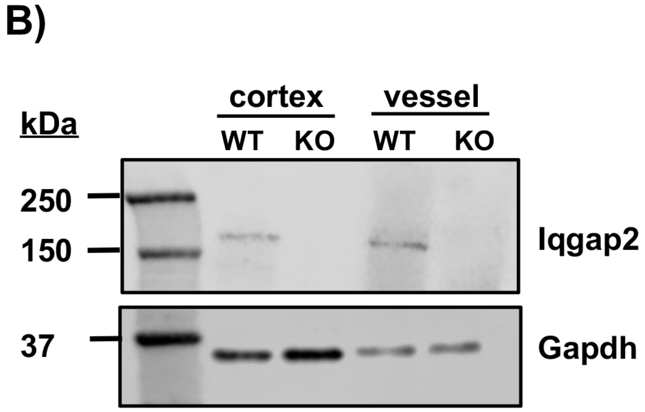

**Supplementary Figure 2: Iqgap2 expression in whole mouse cortex and enriched vessels.**

A) Representative image of isolated brain microvessels stained with lectin.

B) Western blot images of Iqgap2 protein expression in whole cortex and lectin+ vessel fractions isolated from wildtype (WT) and Iqgap2$^{-/-}$ (KO) mice. A single sample was analyzed from 3 pooled cortices or vessel preparations.

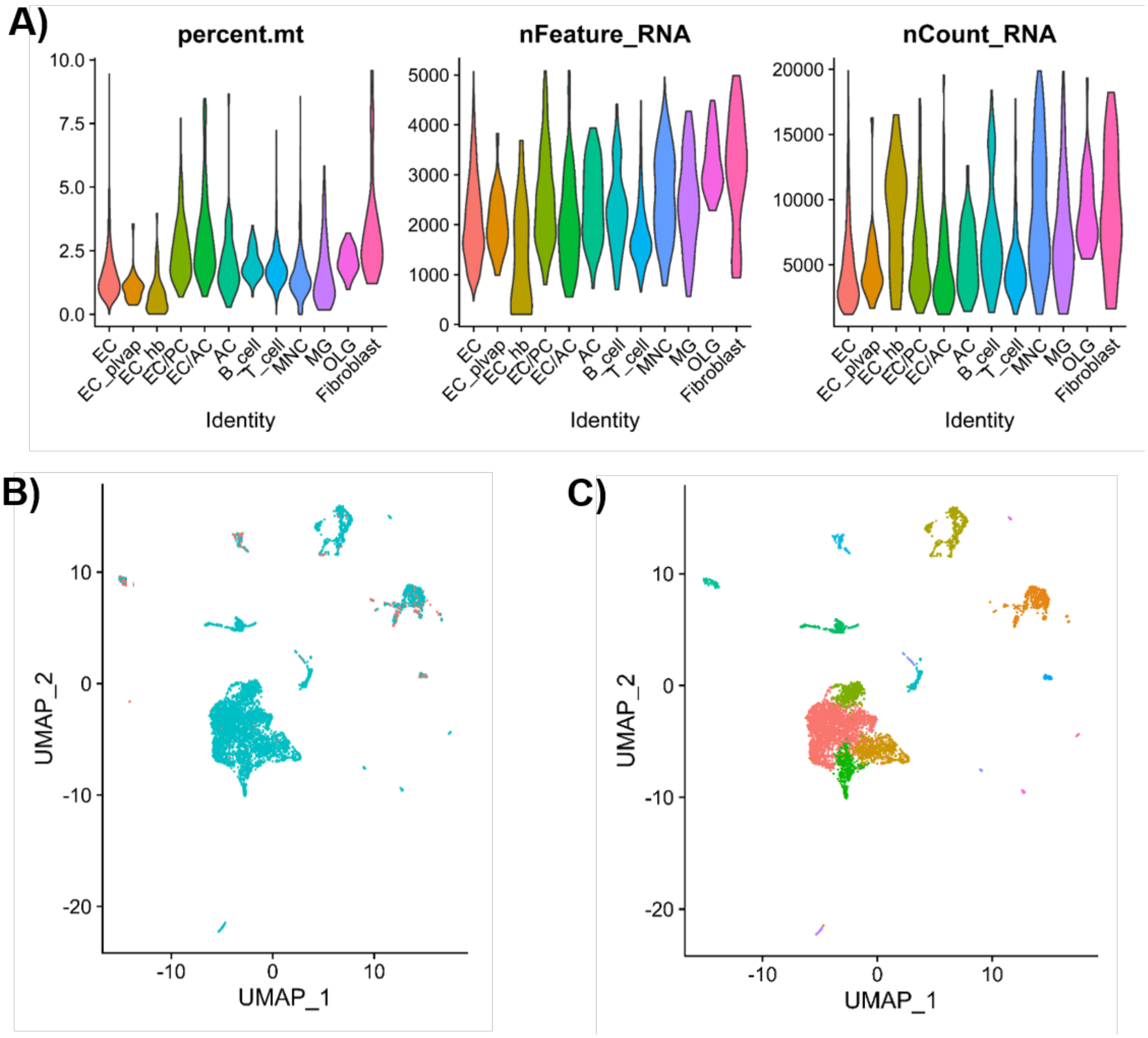

A)
percent.mt
10.0
7.5
5.0
2.5
0.0
nFeature_RNA
5000
4000
3000
2000
1000
0
nCount_RNA
20000
15000
10000
5000
EC
EC_plvap
EC_hb
EC/PC
EC/AC
AC
B_cell
T_cell
MNC
MG
OLG
Fibroblast
Identity
B)
C)
UMAP_2
UMAP_1
10
0
-10
-20

D)

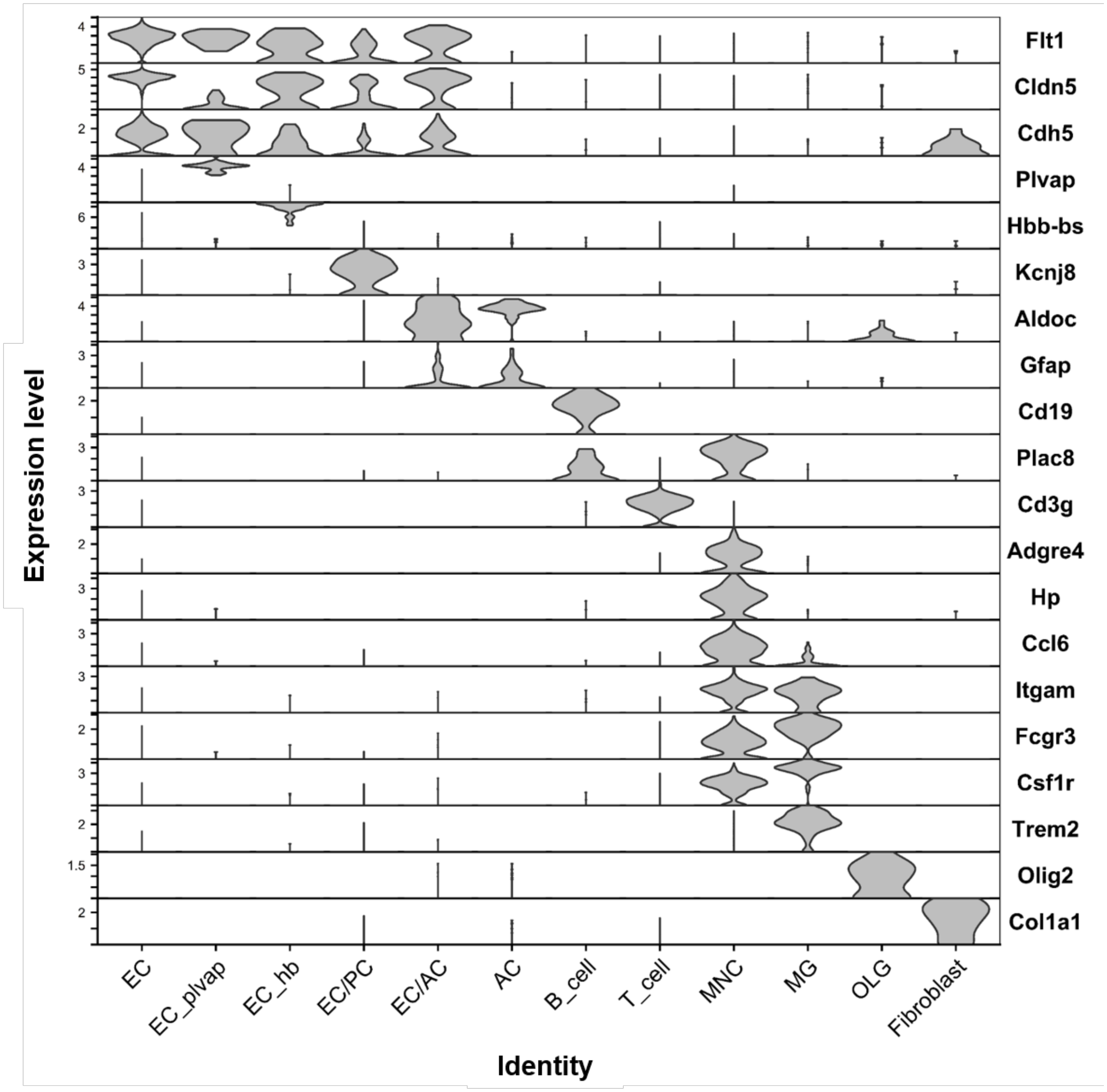

Expression level
Flt1
Cldn5
Cdh5
Plvap
Hbb-bs
Kcnj8
Aldoc
Gfap
Cd19
Plac8
Cd3g
Adgre4
Hp
Ccl6
Itgam
Fcgr3
Csf1r
Trem2
Olig2
Col1a1
EC
EC_plvap
EC_hb
EC/PC
EC/AC
AC
B_cell
T_cell
MNC
MG
OLG
Fibroblast
Identity

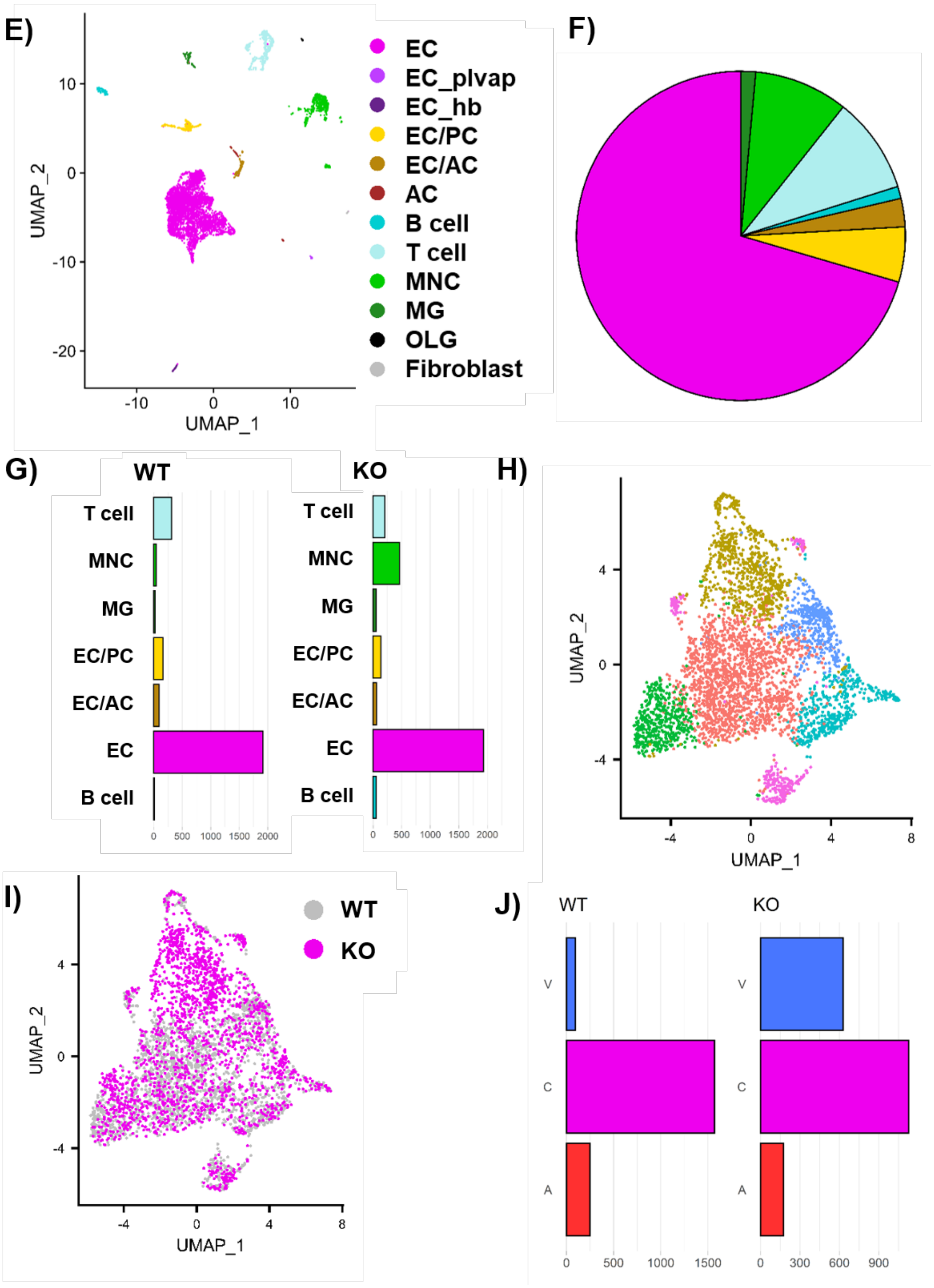

E)
UMAP_2
UMAP_1
10
0
-10
-20
-10
0
10
EC
EC_plvap
EC_hb
EC/PC
EC/AC
AC
B cell
T cell
MNC
MG
OLG
Fibroblast
F)
G)
WT
KO
T cell
MNC
MG
EC/PC
EC/AC
EC
B cell
0
500
1000
1500
2000
H)
UMAP_2
UMAP_1
4
0
-4
-4
0
4
8
I)
WT
KO
UMAP_2
UMAP_1
J)
WT
KO
V
C
A
0
500
1000
1500
0
300
600
900

**Supplementary Figure 3: Expression of cell-type specific marker genes in scRNA-seq samples.**

A) Violin plots displaying percent mitochondrial genes (percent.mt), number of genes per cell (nFeature_RNA) and number of total molecules detected within a cell (nCount_RNA).

B) UMAP of doublet detection analysis. Singlets are labelled in blue and doublets in red.

C) UMAP of unsupervised clustering of all analyzed cells. Colors represent different subclusters.

D) Violin plots of top cell-type specific genes. Cell types are annotated based on expression patterns of these marker genes.

E) UMAP of all annotated clusters based on cell-type specific marker genes. EC=endothelial cells, EC_plvap=PLVAP expressing EC, EC_hb=Hemoglobin expressing EC, EC/PC=EC/stromal cells (pericytes), EC/AC=EC/stromal cells (astrocytes), MNC=monocytes, MG=microglia, OLG=oligodendrocytes.

F) Venn diagram representing cell number contribution of each annotated cluster.

G) Cell numbers for each cell type within WT and KO samples.

H) UMAP of unsupervised clustering of EC cluster.

I) UMAP of genotype origins or cells in the EC cluster.

J) Cell numbers for each zonal identity within the EC cluster.

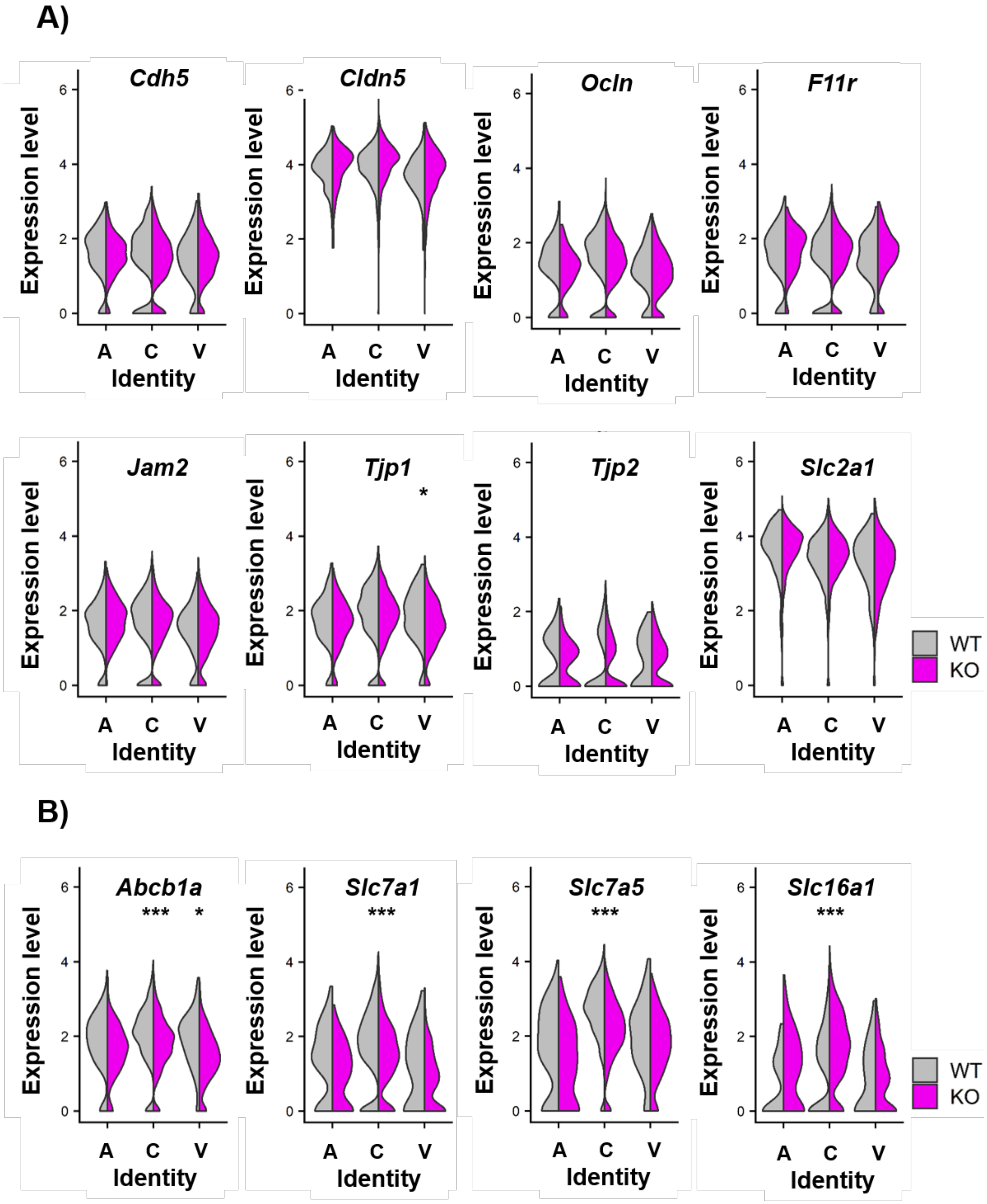
A)
Cdh5
Cldn5
Ocln
F11r
Jam2
Tjp1
*
Tjp2
Slc2a1
Expression level
Identity
A
C
V
WT
KO
B)
Abcb1a
*** *
Slc7a1
***
Slc7a5
***
Slc16a1
***
Expression level
Identity
A
C
V
WT
KO

C)

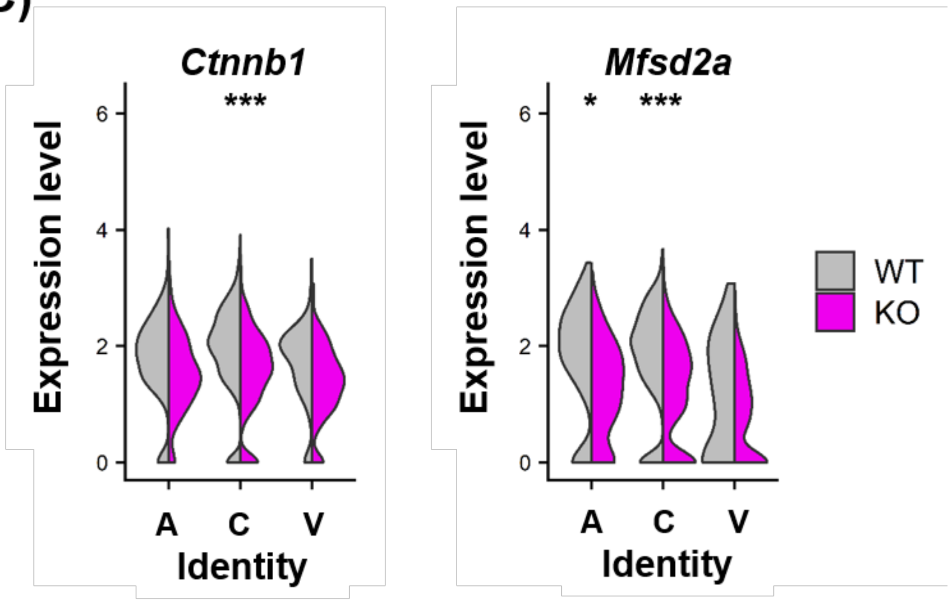

**Supplementary Figure 4: Influence of global Iqgap2 loss on select BBB-relevant genes in BECs.**

A) Split violin plots comparing expression levels of genes encoding junction proteins and Glut1 transporter across vascular zones in WT and KO BECs.

B) Split violin plots comparing expression levels of genes encoding transporter proteins across vascular zones in WT and KO BECs.

C) Split violin plots comparing expression levels of canonical BBB genes across vascular zones in WT and KO BECs.

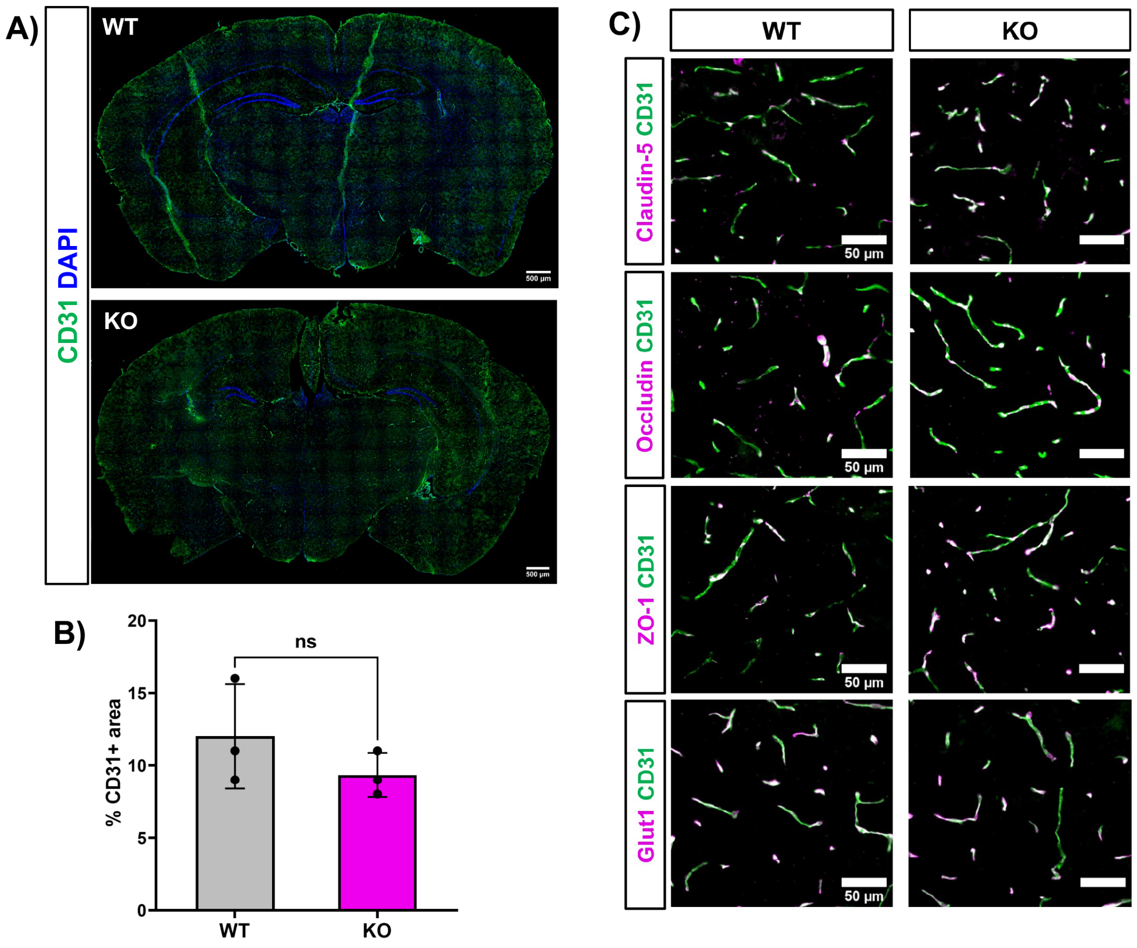

**Supplementary Figure 5: Influence of global Iqgap2 loss on vascular density, tight junction protein expression, and Glut1 protein expression in the brain.**

A) Representative images of brain vasculature in WT and KO mice.

B) Quantification of % CD31+ area. N=3 WT mice and 3 KO mice. Data represented as mean ± SD. Statistical significance was calculated using the student's unpaired t-test.

C) Representative images of vascular claudin-5, occludin, ZO-1, and Glut1 expression in WT and KO mice. Expression patterns were confirmed across N=3 WT mice and 3 KO mice.

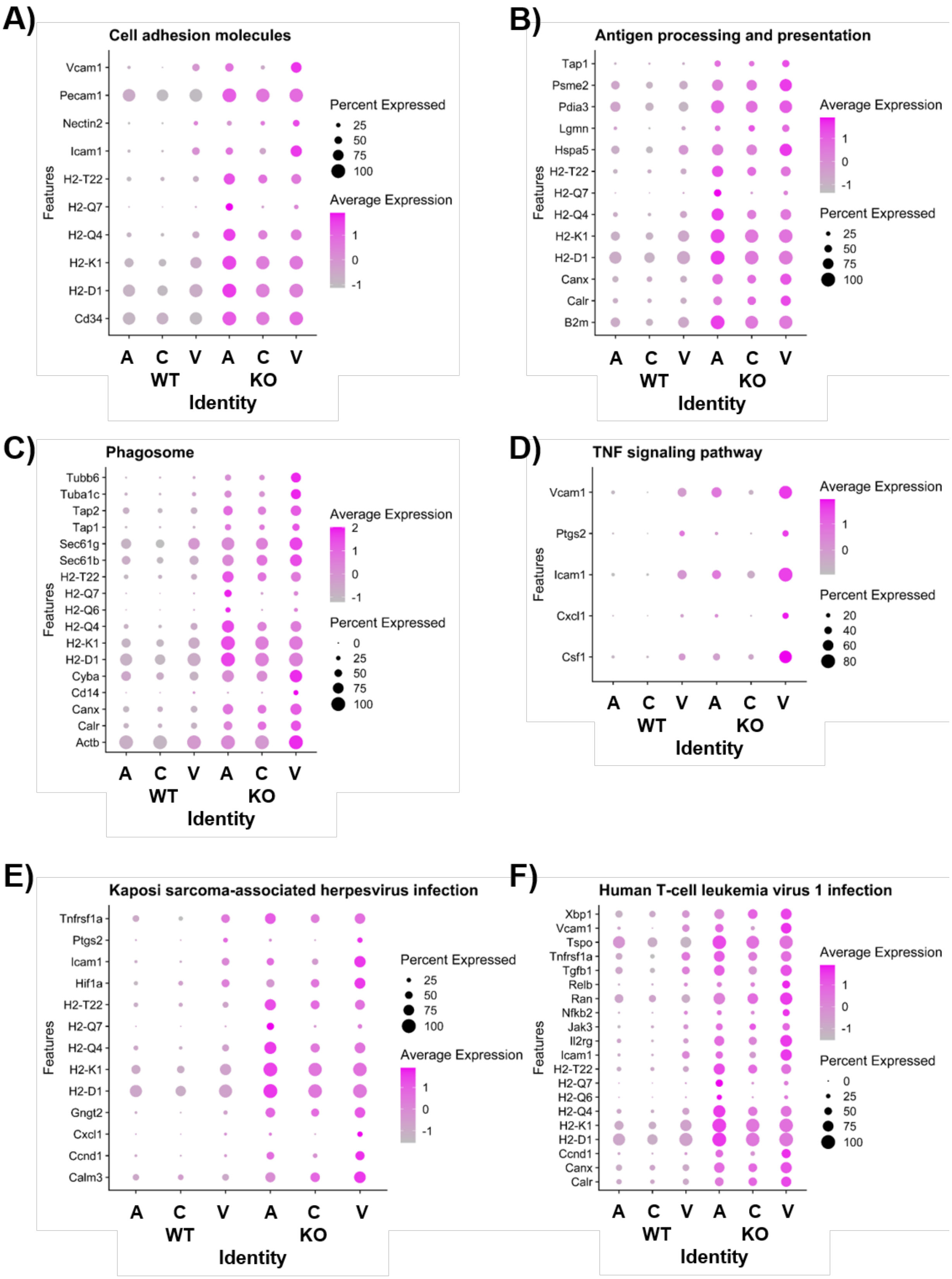
A)
Cell adhesion molecules
Vcam1
Pecam1
Nectin2
Icam1
H2-T22
H2-Q7
H2-Q4
H2-K1
H2-D1
Cd34
Features
Percent Expressed
25
50
75
100
Average Expression
1
0
-1
A C V A C V
WT KO
Identity
B)
Antigen processing and presentation
Tap1
Psme2
Pdia3
Lgmn
Hspa5
H2-T22
H2-Q7
H2-Q4
H2-K1
H2-D1
Canx
Calr
B2m
Features
Average Expression
1
0
-1
Percent Expressed
25
50
75
100
A C V A C V
WT KO
Identity
C)
Phagosome
Tubb6
Tuba1c
Tap2
Tap1
Sec61g
Sec61b
H2-T22
H2-Q7
H2-Q6
H2-Q4
H2-K1
H2-D1
Cyba
Cd14
Canx
Calr
Actb
Features
Average Expression
2
1
0
-1
Percent Expressed
0
25
50
75
100
A C V A C V
WT KO
Identity
D)
TNF signaling pathway
Vcam1
Ptgs2
Icam1
Cxcl1
Csf1
Features
Average Expression
1
0
Percent Expressed
20
40
60
80
A C V A C V
WT KO
Identity
E)
Kaposi sarcoma-associated herpesvirus infection
Tnfrsf1a
Ptgs2
Icam1
Hif1a
H2-T22
H2-Q7
H2-Q4
H2-K1
H2-D1
Gngt2
Cxcl1
Ccnd1
Calm3
Features
Percent Expressed
25
50
75
100
Average Expression
1
0
-1
A C V A C V
WT KO
Identity
F)
Human T-cell leukemia virus 1 infection
Xbp1
Vcam1
Tspo
Tnfrsf1a
Tgfb1
Relb
Ran
Nfkb2
Jak3
Il2rg
Icam1
H2-T22
H2-Q7
H2-Q6
H2-Q4
H2-K1
H2-D1
Ccnd1
Canx
Calr
Features
Average Expression
1
0
-1
Percent Expressed
0
25
50
75
100
A C V A C V
WT KO
Identity

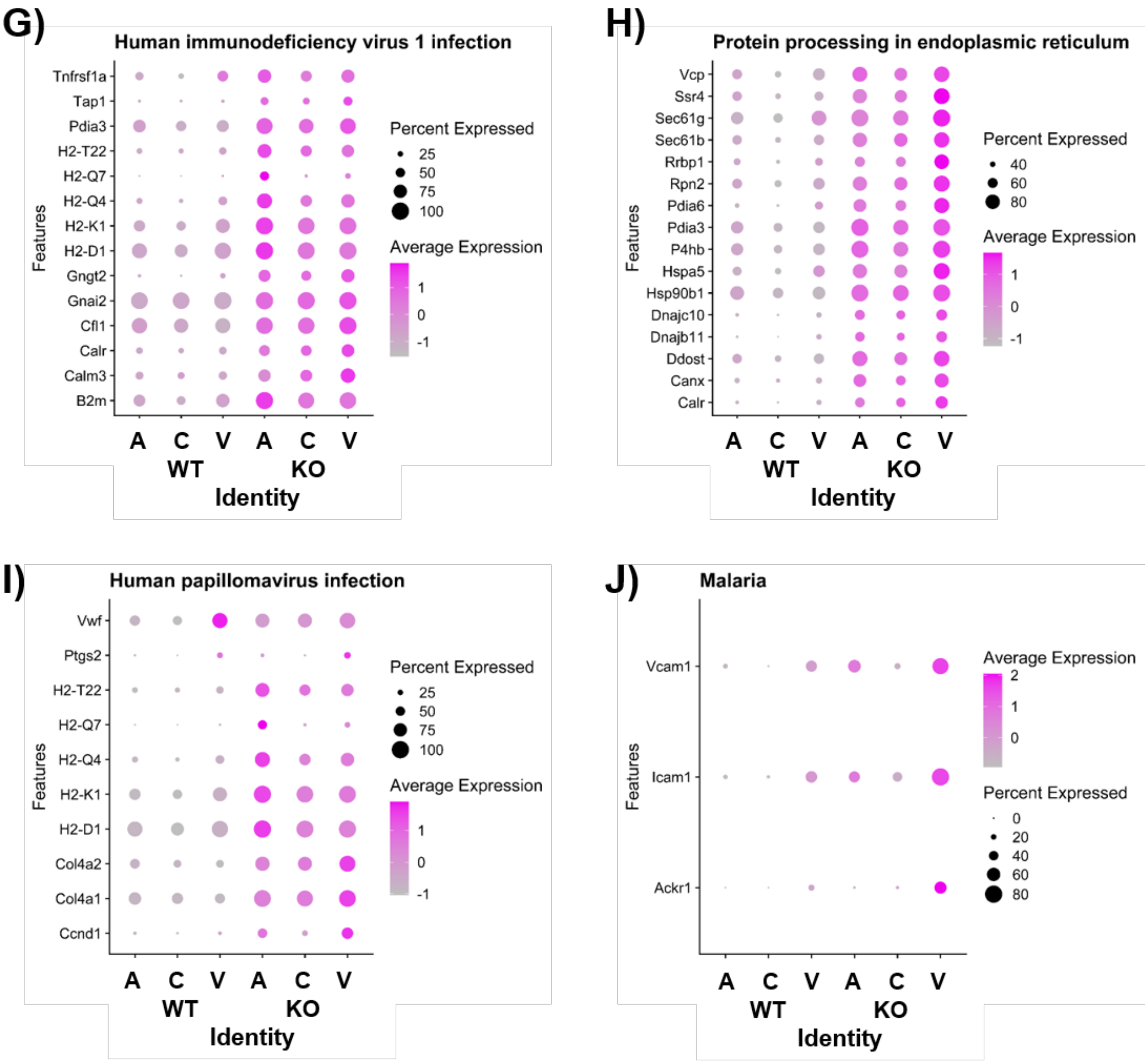

**Supplementary Figure 6: Gene expression changes in zonated BECs corresponding to pathways identified by GSEA.**

In all plots the size of each dot represented percent total cells that express the gene and color represents average expression. The header for each plot across panels A-J indicates the pathway of interest.

A)

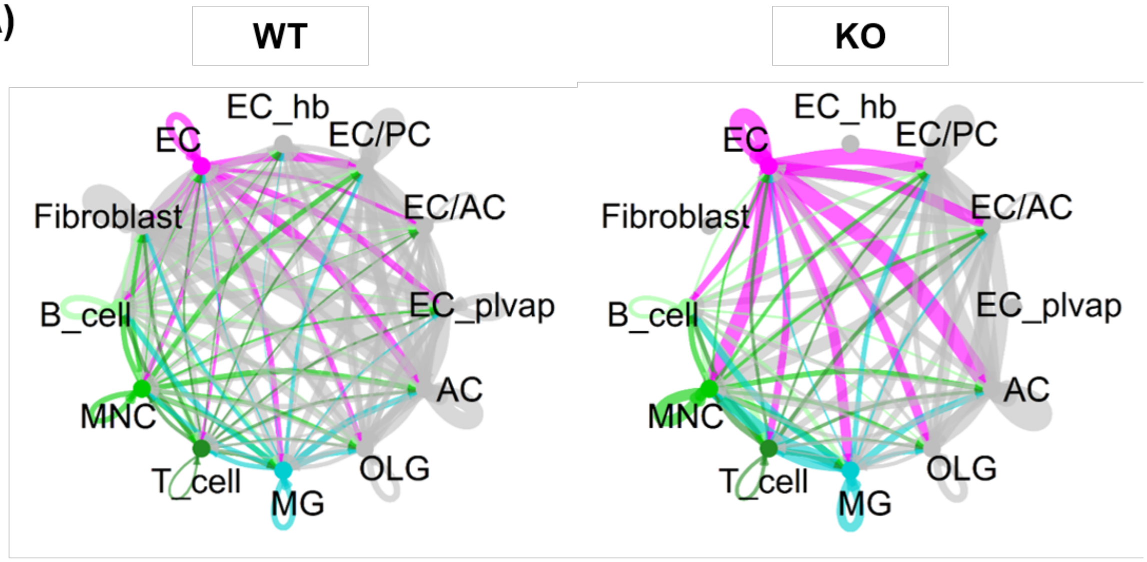
WT
KO
EC_hb
EC
EC/PC
EC/AC
Fibroblast
B_cell
EC_plvap
AC
MNC
T_cell
MG
OLG

B)

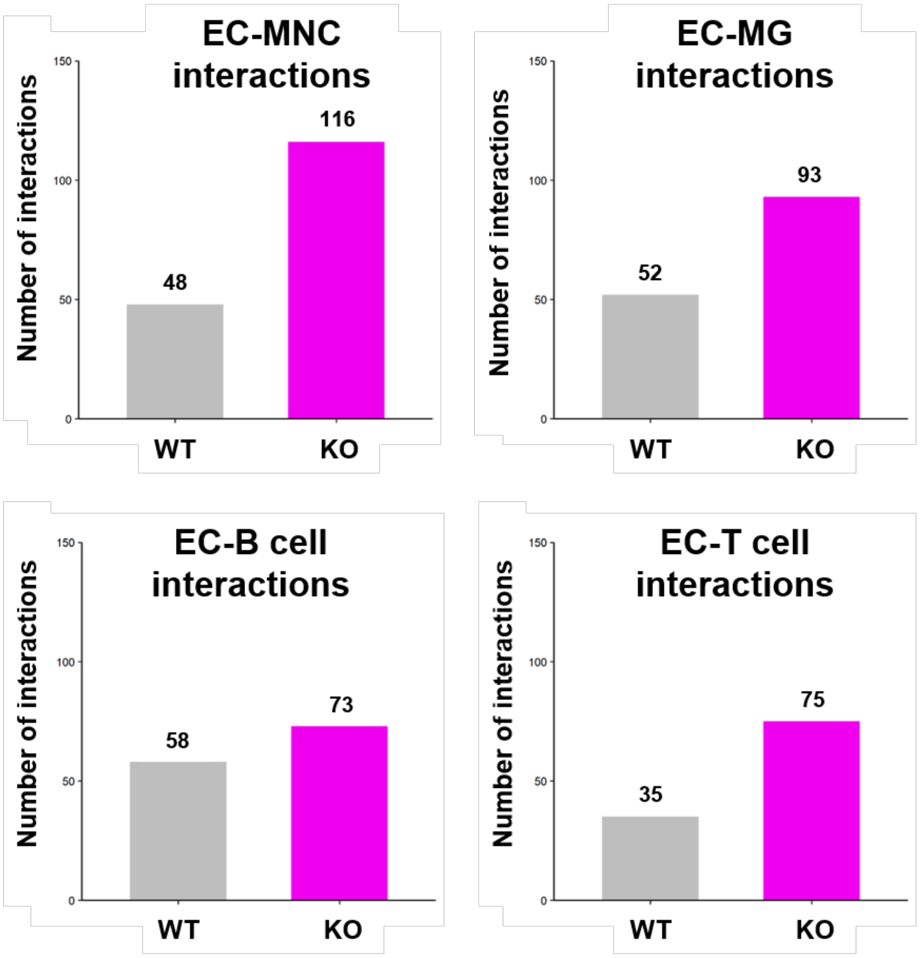
EC-MNC interactions
Number of interactions
150
100
50
0
48
116
WT
KO
EC-MG interactions
Number of interactions
150
100
50
0
52
93
WT
KO
EC-B cell interactions
Number of interactions
150
100
50
0
58
73
WT
KO
EC-T cell interactions
Number of interactions
150
100
50
0
35
75
WT
KO

C)

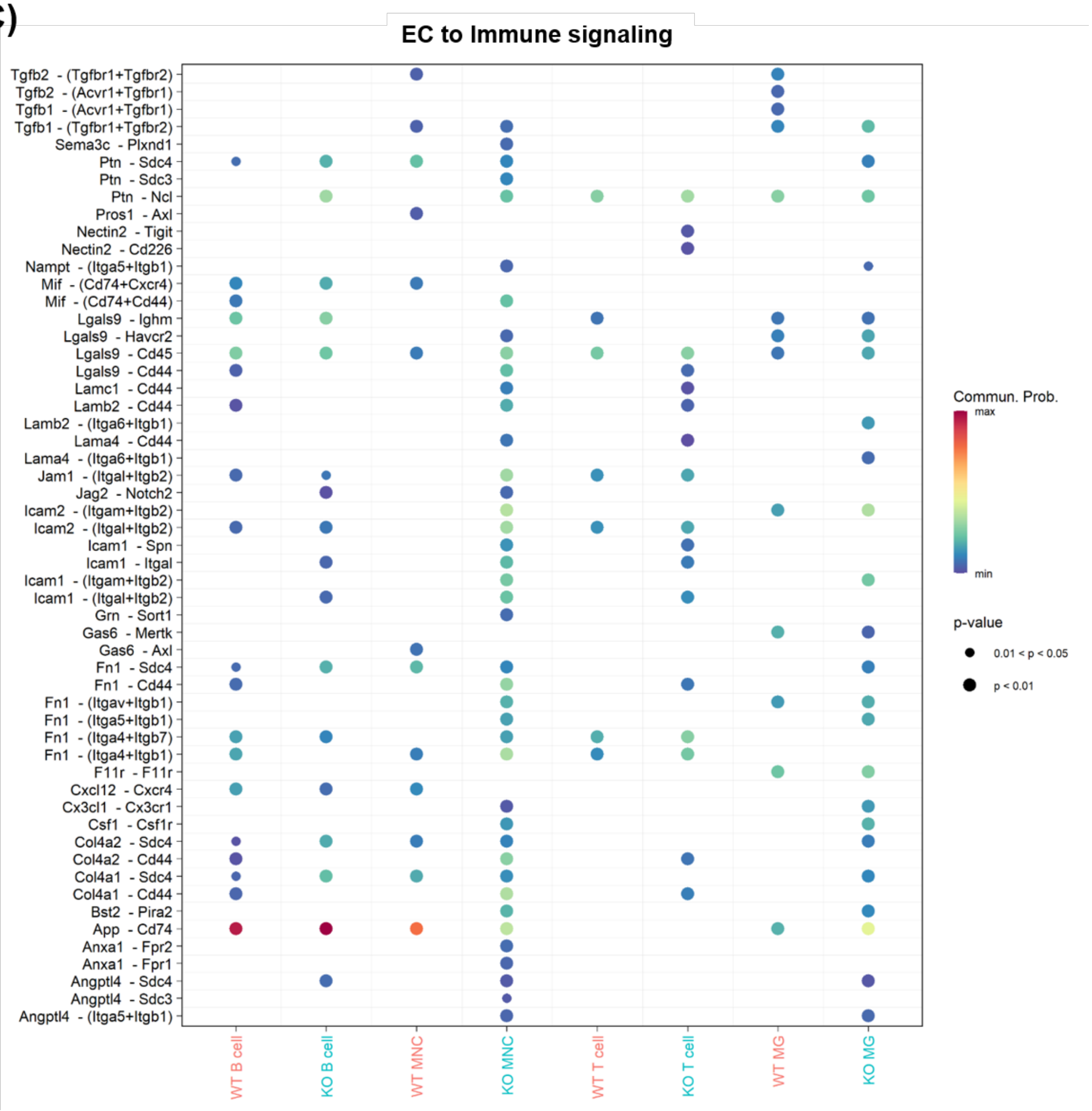
EC to Immune signaling
Tgfb2 - (Tgfbr1+Tgfbr2)
Tgfb2 - (Acvr1+Tgfbr1)
Tgfb1 - (Acvr1+Tgfbr1)
Tgfb1 - (Tgfbr1+Tgfbr2)
Sema3c - Plxnd1
Ptn - Sdc4
Ptn - Sdc3
Ptn - Ncl
Pros1 - Axl
Nectin2 - Tigit
Nectin2 - Cd226
Nampt - (Itga5+Itgb1)
Mif - (Cd74+Cxcr4)
Mif - (Cd74+Cd44)
Lgals9 - Ighm
Lgals9 - Havcr2
Lgals9 - Cd45
Lgals9 - Cd44
Lamc1 - Cd44
Lamb2 - Cd44
Lamb2 - (Itga6+Itgb1)
Lama4 - Cd44
Lama4 - (Itga6+Itgb1)
Jam1 - (Itgal+Itgb2)
Jag2 - Notch2
Icam2 - (Itgam+Itgb2)
Icam2 - (Itgal+Itgb2)
Icam1 - Spn
Icam1 - Itgal
Icam1 - (Itgam+Itgb2)
Icam1 - (Itgal+Itgb2)
Grn - Sort1
Gas6 - Mertk
Gas6 - Axl
Fn1 - Sdc4
Fn1 - Cd44
Fn1 - (Itgav+Itgb1)
Fn1 - (Itga5+Itgb1)
Fn1 - (Itga4+Itgb7)
Fn1 - (Itga4+Itgb1)
F11r - F11r
Cxcl12 - Cxcr4
Cx3cl1 - Cx3cr1
Csf1 - Csf1r
Col4a2 - Sdc4
Col4a2 - Cd44
Col4a1 - Sdc4
Col4a1 - Cd44
Bst2 - Pira2
App - Cd74
Anxa1 - Fpr2
Anxa1 - Fpr1
Angptl4 - Sdc4
Angptl4 - Sdc3
Angptl4 - (Itga5+Itgb1)
WT B cell
KO B cell
WT MNC
KO MNC
WT T cell
KO T cell
WT MG
KO MG
Commun. Prob.
max
min
p-value
0.01 < p < 0.05
p < 0.01

D)

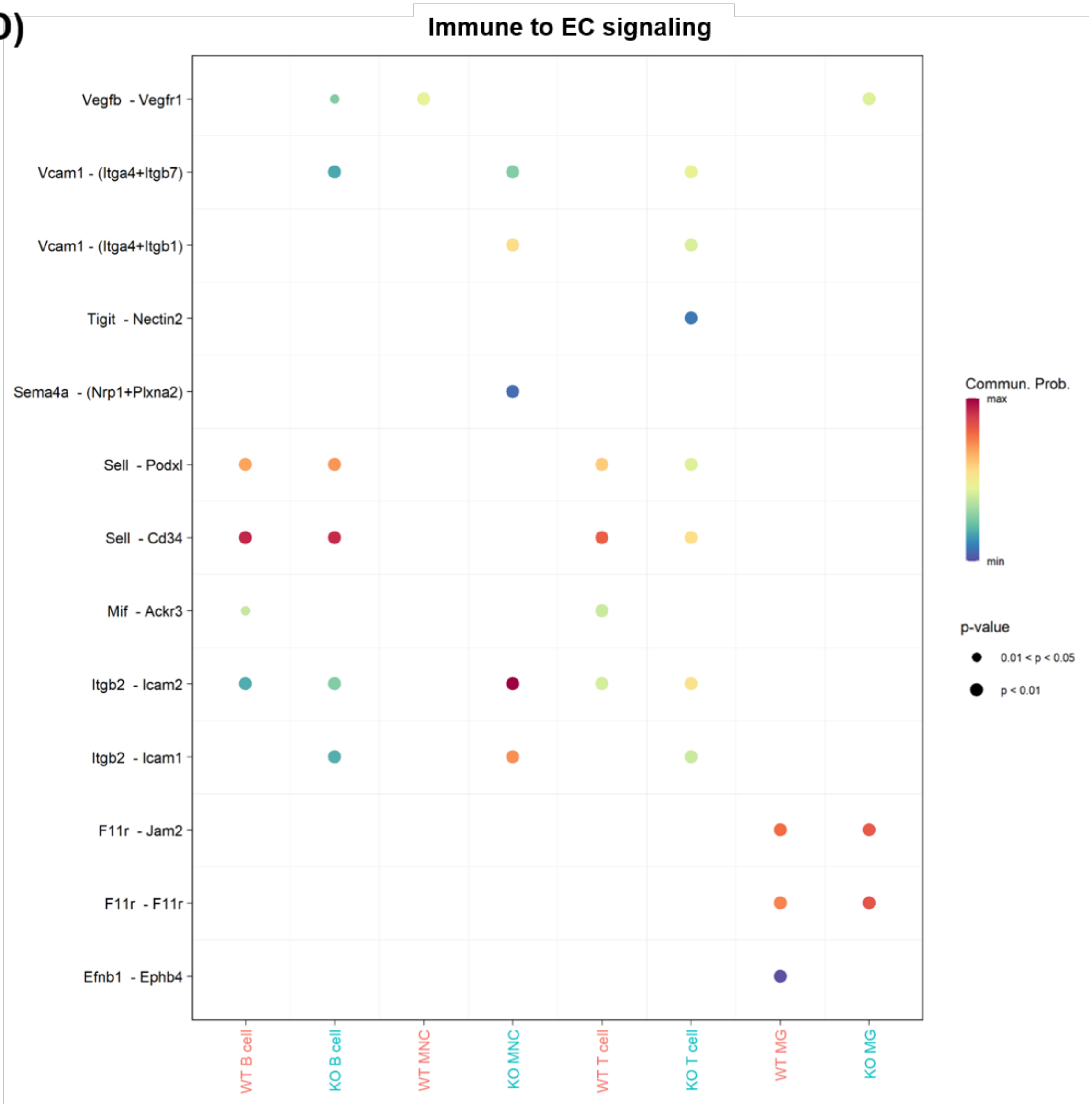

**Supplementary Figure 7: Cell-cell communication between BECs and immune cell types analyzed using CellChat.**

A) Circle plots visualizing cellular communication in WT and KO mice. EC and immune cell interactions are colored as EC (pink), peripheral immune cell types (green), and microglia (blue).

B) Bar charts quantifying EC interactions with microglia (MG), monocytes (MNC), T cells, and B cells.

C) Dot plots indicate predicted EC to immune cell signaling. Colors indicate communication probability and dot size indicates p-value. Statistical significance was calculated using a one-sided permutation test.

D) Dot plots indicate predicted immune cell to EC signaling. Colors indicate communication probability and dot size indicates p-value. Statistical significance was calculated using a one-sided permutation test.

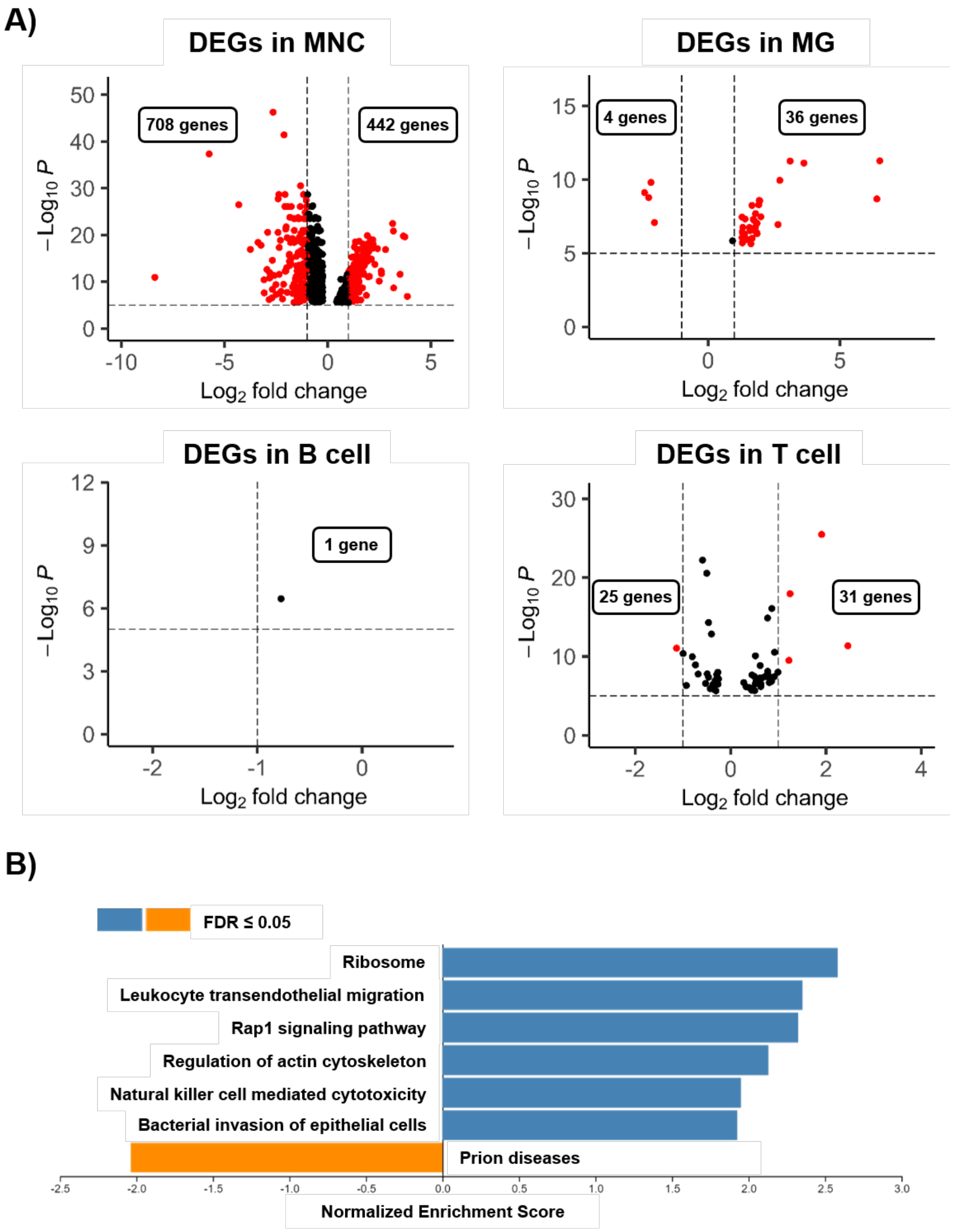

**Supplementary Figure 8: Iqgap2 regulates transcriptomic signatures in immune cells.**

A) Volcano plots show differential gene expression in KO vs WT immune populations: Monocytes (MNC), microglia (MG), B cells, and T cells. All represented dots are significantly altered genes. Red dots represent genes with expression changes by more than 2-fold.

B) KEGG pathway analysis of monocytes shows pathways corresponding to up and downregulated genes.

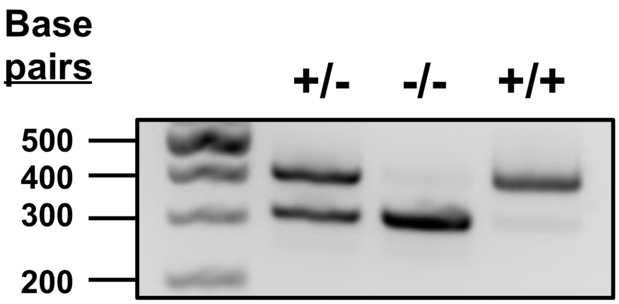

**Supplementary Figure 9: Gel electrophoresis assessment of *Iqgap2* genotype.**

Predicted PCR band sizes are 300 and 391 bp for heterozygotes (+/-), 300 bp for homozygous knockout (-/-), and 391 bp for wild-type animals (+/+).

**Supplementary Table 1: List of differentially expressed genes in brain endothelial cells from wildtype and Iqgap2 null mice segregated by vascular zone.** Table can be found in an accompanying Excel file.

**Supplementary Table 2: Patient sample demographics and de-identified neuropathological information.**

| Case | Age at death/ Sex | Primary clinical categorization | AD neuro-pathology. Braak & Braak | Other neuro-pathologies | PMI | Cause of death | Major comorbidities |
|---|---|---|---|---|---|---|---|
| AD, n1 | 68/F | Alzheimer's disease | Severe (A3, B3, C3) | Mild CAA | 15h | Dementia | Depression |
| AD, n2 | 70/M | Alzheimer's disease | Severe (A3, B3, C3) | Severe CAA | 11h | Dementia | None |
| AD, n3 | 81/M | Alzheimer's disease | Mild (A2, B1, C2) | Severe CAA, 3cm hemorrhage | 19h | Dementia/ seizure | Atrial fibrillation |
| AD, n4 | 81/F | Alzheimer's disease | Severe (A3, B3, C3) | Severe CAA | 36h | Dementia | Athero-sclerotic heart disease |
| AD, n5 | 61/M | Alzheimer's disease | Moderate-Severe (A2, B2, C3) | Moderate-Severe CAA | 10h | Dementia | Intracerebral brain hemorrhage |
| AD, n6 | 68/F | Alzheimer's disease | Severe (A3, B3, C3) | Mild CAA | 9h | Dementia | Mild cognitive impairment |
| AD, n7 | 66/F | Alzheimer's disease | Severe (B5, Thal 4, C3) | Mild CAA | 25h | Dementia | Hypertension |
| non-AD, n1 | 66/F | Neurological control | None | None | 33h | Leukemia | COPD, liver transplant recipient |
| non-AD, n2 | 72/M | Neurological control | None | None | 12h | Acute respiratory distress syndrome | Renal failure, heart failure. |
| non-AD, n3 | 87/M | Non-AD | None | Stroke | 6h | Stroke | Arteriolo-sclerosis |
| non-AD, n4 | 87/M | Non-AD | Rare amyloid plaques | None | 13h | Myocardial infarction | Coronary artery disease and heart failure |
| non-AD, n5 | 66/M | Non-AD | None | None | 24h | Septic shock | Diabetes, renal transplant |
| non-AD, n6 | 70/F | Non-AD | None | None | 35h | Cardiovascular decease | Arteriolo-sclerosis |
| non-AD, n7 | 71/M | Non-AD | Moderate vascular disease | Low-grade dementia | 29h | Cardiovascular decease | Hypertension |

**Supplementary Table 3: Primary antibodies.**

| Target | Vendor | Product Number | Species | Dilution | Use |
|---|---|---|---|---|---|
| Collagen | DSHB | M3F5 | Mouse | 1:100 | IHC |
| IQGAP2 | - | - | Rabbit | 1:50 | IHC |
| Iqgap2 | Santa Cruz Biotechnology | sc-55525 | Mouse | 1:500 | WB |
| GAPDH (D16H11) | Cell Signaling Technologies | 5174 | Rabbit | 1:1000 | WB |
| Vcam1 | Santa Cruz | sc-13160 | Mouse | 1:500 | IHC |
| CD31 (MEC 13.3) | BD Biosciences | 553370 | Rat | 1:200 | IHC |
| CD45 (30-F11) | BD Biosciences | 550539 | Rat | 1:100 | IHC |
| Claudin-5-488 | Invitrogen | 352588 | - | 1:100 | IHC |
| Occludin-594 | Invitrogen | 331594 | - | 1:100 | IHC |
| ZO1-594 | Invitrogen | 339194 | - | 1:100 | IHC |
| GLUT1-488 | Abcam | ab195359 | - | 1:300 | IHC |

**Supplementary Table 4: Secondary antibodies and other detection reagents.**

| Species Reactivity | Vendor | Product Number | Host | Conjugate | Dilution | Use |
|---|---|---|---|---|---|---|
| Rabbit | LI-COR | 926-32211 | Goat | IRDye 800 | 1:15000 | WB |
| Rat | Invitrogen | A21247 | Goat | AlexaFluor 647 | 1:500 | IHC |
| Rat | Invitrogen | A11006 | Goat | AlexaFluor 488 | 1:500 | IHC |
| Rabbit | Invitrogen | A21206 | Donkey | AlexaFluor 488 | 1:500 | IHC |
| Rabbit | Invitrogen | A31572 | Donkey | AlexaFluor 555 | 1:500 | IHC |
| Mouse | Invitrogen | A21202 | Donkey | AlexaFluor 488 | 1:500 | IHC |
| Rabbit | Abcam | ab150073 | Donkey | AlexaFluor 488 | 1:1000 | IHC |
| Mouse | Abcam | ab150108 | Donkey | AlexaFluor 594 | 1:1000 | IHC |
| - | Vector Labs | DL-1174-1 | Lectin | DyLight 488 | 1:100 | IHC |
| - | Vector Labs | DL-1178-1 | Lectin | DyLight 649 | 1:100 | IHC |
